# Walking strides direct rapid and flexible recruitment of visual circuits for course control in *Drosophila*

**DOI:** 10.1101/2021.10.10.463817

**Authors:** Terufumi Fujiwara, Margarida Brotas, M Eugenia Chiappe

## Abstract

Flexible mapping between activity in sensory systems and movement parameters is a hallmark of successful motor control. This flexibility depends on continuous comparison of short-term postural dynamics and the longer-term goals of an animal, thereby necessitating neural mechanisms that can operate across multiple timescales. To understand how such body-brain interactions emerge to control movement across timescales, we performed whole-cell patch recordings from visual neurons involved in course control in *Drosophila*. We demonstrate that the activity of leg mechanosensory cells, propagating via specific ascending neurons, is critical to provide a clock signal to the visual circuit for stride-by-stride steering adjustments and, at longer timescales, information on speed-associated motor context to flexibly recruit visual circuits for course control. Thus, our data reveal a stride-based mechanism for the control of high-performance walking operating at multiple timescales. We propose that this mechanism functions as a general basis for adaptive control of locomotion.

## Introduction

Adaptive behavior—behavior that enhances survival in complex natural environments—depends on the capacity of the central nervous system to flexibly engage functional networks that control movement parameters ^1–5^. This flexibility operates according to internal contexts that are defined by continuous feedback on how behavioral goals match the state of the body (i.e., motor context), the animal’s longer-term physiological needs (i.e., internal states), and its previous experience ^6^. Thus, the contextdependent recruitment of circuits for movement control occurs by combining diverse streams of information across different timescales ^7^. Work on context-dependent modulation of brain circuits has largely focused on the effect of internal states ^8–10^ and experience ^11–16^. However, much less is known about how the central nervous system is organized to signal motor context and orchestrate the flexible recruitment of brain circuits for the control of movement (**Fig. 1A**). This lack of fundamental knowledge represents a major limitation in efforts to understand the neural mechanisms underlying adaptive behavior.

**Figure 1:**
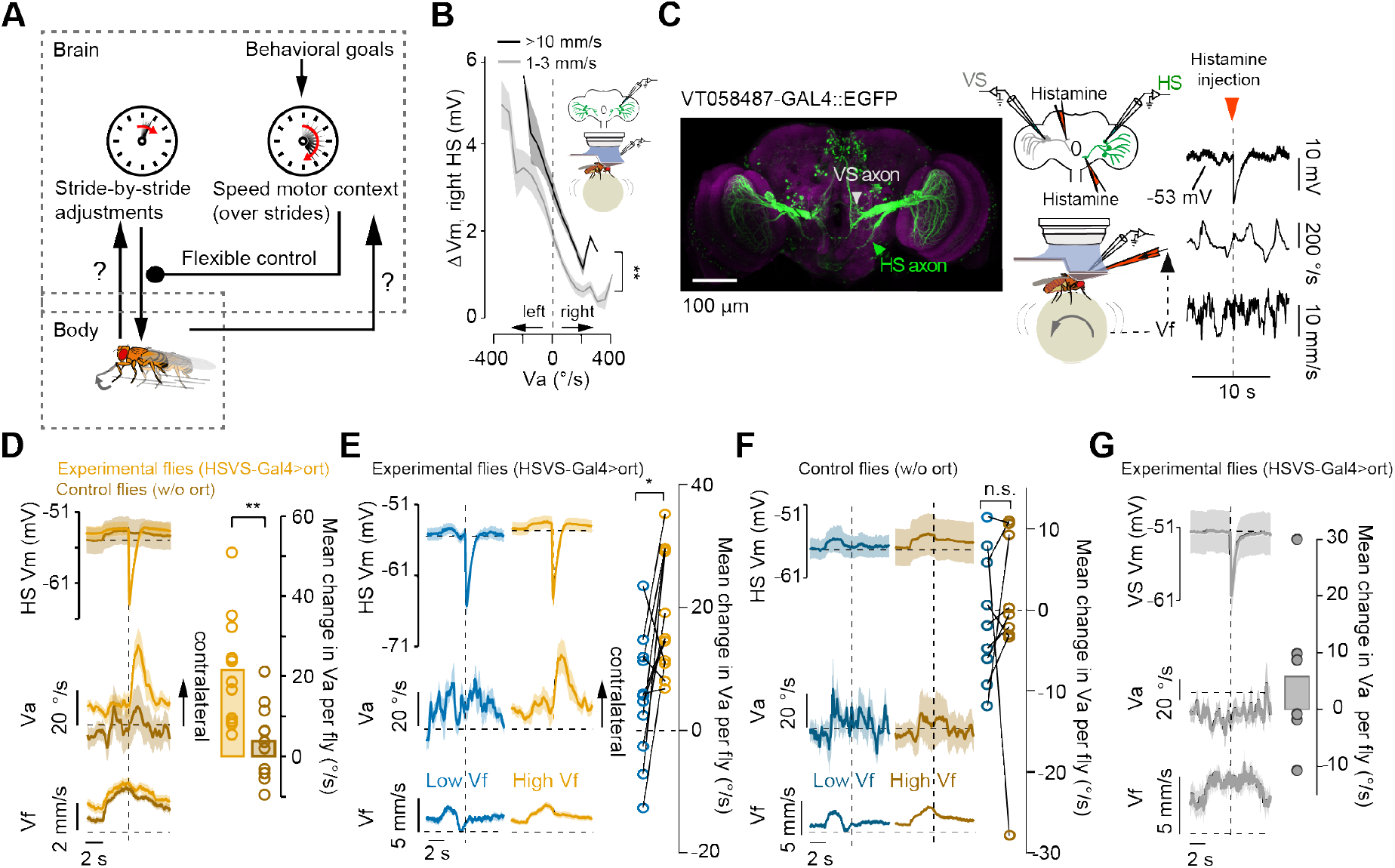
HS cells contribute to steering in high-but not low-speed walking bouts. **(A)** Walking performance depends on the continuous comparison between short-term postural dynamics and the longer-term behavioral goals of an animal. This comparison defines a motor context that flexibly recruits brain circuits for rapid, goal-directed movement adjustments. The mechanisms underlying such control are unknown; we propose that signals from the ventral nerve cord may convey information about the state of the body at multiple timescales to recruit brain circuits raoidly and flexibly. (B) Simultaneous recordings of neural activity (right HS cells) and walking behavior in darkness (inlet). Change in membrane potential relative to quiescence (ΔVm), as a function of the fly’s angular velocity (Va) in walking bouts with average low (1–3 mm/s, gray) or high (>10 mm/s, black) forward velocity (Vf, grand mean ± SEM, N=9 fly-cell pairs, n=556 walking bouts). **: p=0, for the offset difference between the curves, bootstrapping method (see **Methods). (C)** Left, confocal image stack of the VTO58487 line driving the expression of EGFP. HS- and VS-cell axon terminals are indicated with arrowheads. Middle, simultaneous whole-cell patch recordings from HS or VS cells and walking activity in darkness. A closed-loop system between Vf and the pressure ejection system controlled the conditional application of histamine. Right, example traces of the membrane potential of an HS cell expressing the histamine receptor ort (VT058487>ort), and Va and Vf. The arrowhead indicates a histamine injection at the HS axon’s terminals contingent to a mean high Vf over 3 seconds. (D) Left, Vm, Va, and Vf traces (grand mean ± SEM) in experimental (orange, N=12 flies) or control flies (without ort expression, maroon, N=11 flies) triggered at histamine application. Right, mean change in Va per fly before and after histamine injection in experimental vs. control flies (p=0.0023, Z=3.05, Wilcoxon rank-sum test). Note that Vm is reported for a subset of flies in which the whole-cell condition lasted until the end of the experiment (7/12 and 6/11 fly-cell pairs for experimental and control flies, respectively). (E) Left, Vm, Va, and Vf triggered at histamine injection conditioned to low (blue) vs. high (orange) Vf in experimental flies (grand mean ± SEM). Right, mean change in Va per fly upon histamine injection at low vs. high Vf (right, p=0.019, Wilcoxon signed-rank test, N=11 flies). (F) Same as (E) but for control flies (p=0.36, Wilcoxon signed-rank test, N=9 flies). Note that Vm is only reported for recordings that lasted until the end of the experiment (4/9 fly-cell pairs for control flies). (G) Histamine was injected at the axon terminals of VS-cells while their activity was monitored by whole-cell patch recordings. Vm, Va, and Vf triggered at histamine application (N=6 flies, grand mean ± SEM). Note that Vm is reported only for recordings that lasted until the end of the experiment (5/6 fly-cell pairs). Right, mean change in Va per fly upon histamine injection (before injection: −25.8±6.8 °/s vs. after injection: −20.0±8.1 °/s, p=0.69, Wilcoxon signed-rank test, N=6 flies)

The emergence of an internal motor context likely depends on recurrent interactions between brain premotor centers and the spinal cord, operating across multiple timescales (**Fig. 1A**). Indeed, the spinal cord houses as many ascending fibers as descending ones ^17^, and yet very little is known about the behavioral function of this ascending information ^18–20^. During walking, coupling of ascending modulation in the mammalian brain to individual strides suggests that supraspinal circuits receive immediate information about the state of the body ^21^. However, the exact function and nature of this stride-coupled modulations remain unknown, partly due to the complexity and highly distributed structure of mammalian brain premotor circuits, and to the rather limited understanding of how activity within these circuits leads to different aspects of walking behavior ^22^.

The compact central nervous system of *Drosophila melanogaster* provides a powerful model in which to study the mechanisms and timescales through which motor context-related signals emerge and impact neural processing and walking control. However, activity dynamics in central brain circuits have not been characterized at the level of stride-by-stride timescales during walking. Importantly, and in contrast to internal states, signals related to the state of the body can be directly measured by detailed quantitative analysis of behavior and neural physiology, allowing the nature and mechanisms of motor-context modulation and its effect on movement adjustments to be dissected. In insects, the posterior slope (PS), a premotor region with strong multisensory convergence ^23^, provides output to several populations of descending neurons known to be involved in steering maneuvers during walking (Namiki and Kanzaki, 2016; Rayshubskiy et al., 2020). The PS receives inputs from higher order centers, from the ventral nerve cord (VNC, the insect analogue of the spinal cord) via ascending neurons, and from visual motion pathways ^24–27^. One class of visual neurons projecting to the PS is the genetically identified population of HS cells, which are readily accessible for physiological recording and manipulation during behavior ^28–30^. Similar to some descending neurons ^31,32^, unilateral activation of HS cells promotes ipsilateral body steering during walking and flight ^28,33,34^. The activity of HS cells also correlates with the forward velocity of the fly ^28^, although the function of this signal has remained unclear. One possibility is that a speed-related signal provides information on motor context, either related to the behavioral goals of the fly (“run forward”) or to the current state of the body (“walking at high speed”), or both. Thus, this visuomotor circuit provides an ideal model in which to examine the direct role of motor-context information on the mapping between neural activity dynamics in premotor circuits and specific aspects of walking behavior.

Here, we combine whole-cell patch recordings in walking flies with optogenetics and targeted suppression of chemical synapses to uncover a speed-related context signal originating in leg-mechanosensory neurons that modulates activity in HS cells across multiple timescales via an identified class of ascending neurons (ANs). On a stride-by-stride basis, this signal tunes activity in HS cells for rapid steering maneuvers. Over several similar strides, a slower modulation emerges and recruits HS cells to steering adjustments in a specific motor context that correlates with stance duration. Thus, a single source, the stride, operating at multiple timescales provides an elegant solution to flexibly engage a functional network in movement corrections within a continuous behavior that is rarely in steady state. We propose that these findings represent a general mechanism by which bidirectional interactions between the peripheral nervous system and brain visual circuits contribute to the high-performance control of locomotion.

## Results

### HS cells contribute to steering in a forward velocity-dependent manner

HS cells are thought to contribute to course control under conditions in which the fly actively maintains the direction of locomotion, such as when running forward at high speed. If a forward speed-associated signal in the dynamics of HS cells functions as a contextual signal, two properties should be observed at different walking speeds. First, the selectivity of HS-cell responses should not change as a function of walking speed. Second, manipulating the activity of HS cells should lead to steering effects in a forward velocity-dependent manner. To test the first prediction, we examined the direction-selective properties of HS cells at high vs. low forward velocity by performing whole-cell patch recordings from HS cells in flies walking in darkness. We excluded visual stimulation in these experiments, since its presence influences the walking speed of the fly ^35^. Activity in HS cells was selective for the direction of the fly’s rotations independent of walking speeds. However, the overall activity of the neurons was greater at high than low forward velocity (**Fig. 1B**). Thus, the walking speed of the fly induces a modulatory effect on the level of depolarization of HS cells, likely affecting synaptic activity in these non-spiking neurons. This finding suggests that speed-related modulation may impact the contribution of HS-cell activity to course control.

Previous work showed that balanced activity in the bilateral population of HS cells is important for walking control ^28,33^. Therefore, to evaluate a putative speed-dependent effect of HS-cell activity on behavior, we performed conditional unilateral silencing using a chemogenetic strategy ^36^ (see **Methods**). We expressed the histamine-gated chloride channel *ort* in the bilateral population of HS cells and employed local application of histamine at the axon terminals of HS cells from the right hemisphere to induce a unilateral perturbation. To test the effect of conditional silencing of HS cells on walking behavior, we then controlled the application of histamine according to the average forward velocity (Vf) of the fly for at least 3s (**Fig. 1C,** see **Methods**). Application of histamine led to a prominent inhibition of neural activity in Ort-expressing HS cells independently of the walking speed of the fly (**Fig. 1D, E**). However, the overt effect of this manipulation on behavior was only apparent in flies walking at high Vf. Consistent with previous work ^33^, silencing the right-sided population of HS cells promoted a robust contralateral left rotation in experimental but not control flies at high Vf, whereas no effect was observed at low Vf (**Fig. 1D–F**). Importantly, the effect on behavior was independent of the expression of Ort in other visual neurons also labeled in the transgenic line (VS cells, **Fig. 1C, G**). We conclude that the forward speed-related signal in the dynamics of HS cells represents a motor-context modulation that allows flexible recruitment of HS-cell activity for steering adjustments.

### A forward velocity-related signal modulates activity in HS cells at multiple timescales

In a continuous behavior such as walking, Vf can fluctuate at different timescales. Over several seconds, fluctuations in Vf may reflect changes in motor programs or behavioral goals during several strides (**Fig. 1**), whereas on a shorter, stride-by-stride timescale, changes in Vf reflect leg reactive forces ^37–39^. Variability of Vf at this short timescale may reflect unexpected perturbations in the neuromechanical system within a single stride that must be corrected for in the next stride according to the motor programs and behavioral goals at play (**Fig. 1A**). Thus, if HS cells are recruited for adjustments at such short timescales, their modulation of activity should not only occur over seconds, reflecting the overall state of Vf, but also at shorter timescales supporting stride-by-stride steering.

In darkness, the fly’s spontaneous behavior was characterized by highly fluctuating Vf and angular velocity (Va), and the activity of HS cells was influenced by both (**Fig. 2A,B**) ^28^. Occasionally, even in darkness, the fly maintained a stable course with very low-amplitude angular movements and high Vf (**Fig. 2B,** gray shadow). Interestingly, mapping the activity of HS cells onto the virtual path of such walking segments revealed fast periodic oscillations as the fly walked forward (**Fig. 2C**), suggesting that Vf-related signals modulate activity in HS cells at short timescales. To test this hypothesis, we examined the correlation between neural activity and Vf at different temporal frequencies. For comparison, we also analyzed the correlation between neural activity and Va. At temporal frequencies characteristic of high walking speed on the ball (>5Hz) ^28^, the correlation between neural activity and walking velocities was dominated by Vf (**Fig. 2D**). In contrast, at lower frequencies (<5Hz), this correlation was dominated by Va. Altogether, these results suggest that HS cells can be modulated by Vf-associated signals at multiple timescales: both at a stride timescale and over the course of a few seconds (**Fig. 1C–F**).

**Figure 2.**
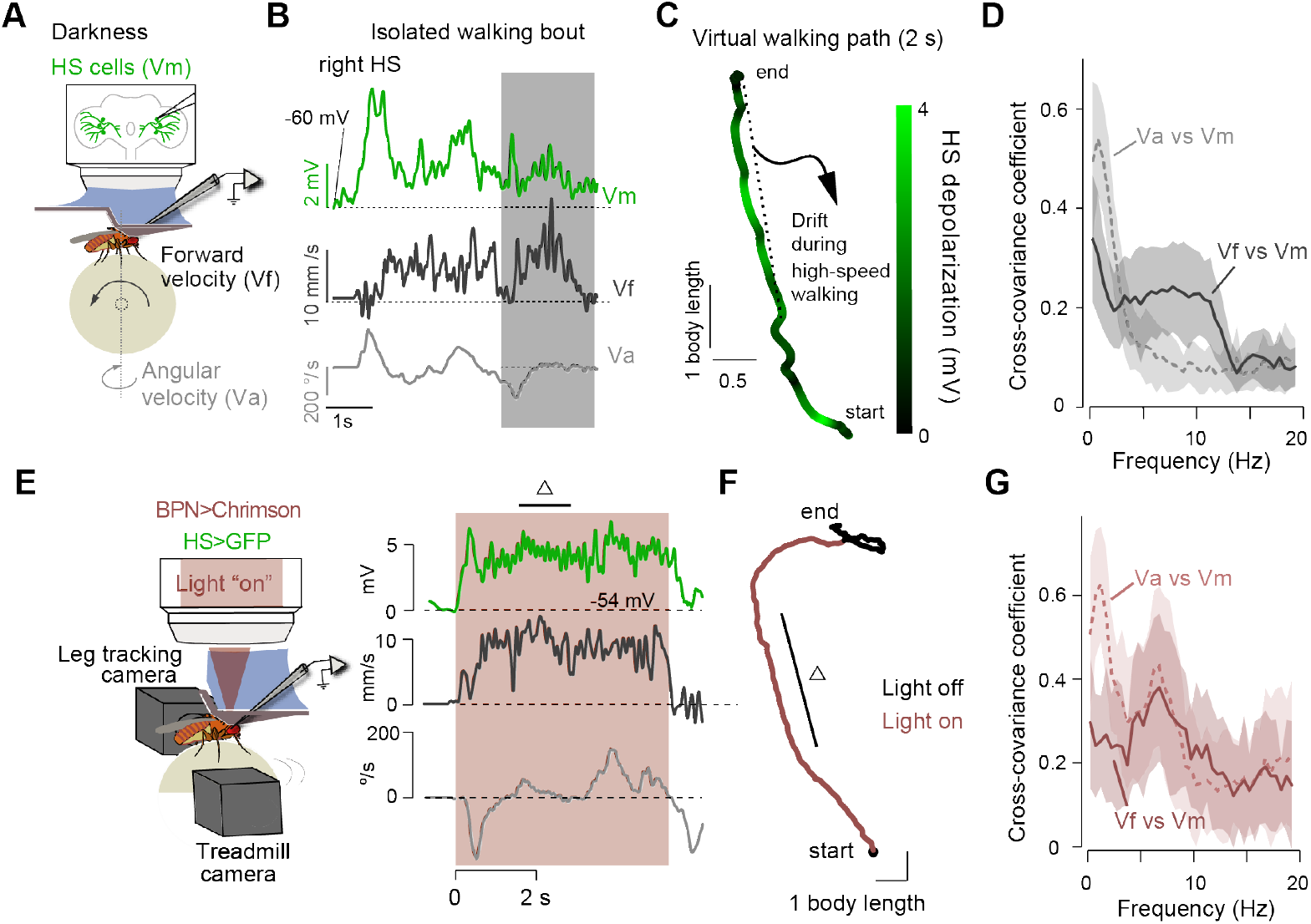
HS cells activity and forward velocity oscillate at high frequencies during spontaneous and induced high-speed walking bouts. **(A)** Whole-cell patch recordings were performed from HS cells in flies walking in darkness. **(B)** Example traces of a right-side HS cell’s membrane potential (Vm), the fly’s forward (Vf, black) and angular (Va, gray) velocities. The shade highlights a moment when the fly walked spontaneously with high forward speed and low angular velocity. (C) A segment of a virtual walking path reconstructed from the treadmill signals during the shaded segment in **(B).** The path is color-coded as a function of the cell depolarization level. **(D)** Maximal coefficient of the cross-covariance between the Vm and Va (gray, dashed line), and between the Vm and Vf (black trace) as a function of the temporal frequency of the respective signals (grand mean ± SEM, n=25 fly-cell pairs). (E) Left, schematic of the experimental setup for tracking neural activity, body velocities and leg movements in simultaneous. Right, example time series of a right HS cell’s Vm, the fly’s Vf (black) and Va (gray) during BPN activation while walking in darkness. The orange shaded area indicates the optogenetic activation period. (F) Reconstructed virtual path of the example trial shown in (E). Walking under BPN activation is indicated by “lights on”. Hie triangles in (E) and (F) highlight a moment when the fly displayed straight walking. **(G)** Same as in **(D)** but for experiments under BPN activation (n=19 fly-cell pairs).

### An optogenetic paradigm to study modulation of HS cells across multiple timescales

To ensure meaningful physiological results, we sought to establish experimental conditions in which flies display frequent walking bouts with high Vf. Because spontaneous walking behavior on the ball in darkness varies substantially across head-fixed individuals, we developed an optogenetics-based paradigm to promote walking bouts at high speed more consistently (**Fig. 2E, F**). To achieve this, we expressed the light-gated cation channel CsChrimson ^40^ in bolt protocerebral neurons (BPNs), a recently identified population of interneurons whose activity induces forward runs in flies ^31^. Light activation of BPNs induced high Vf walking bouts that were accompanied by low Va (absolute mean±SEM=73.6±3.0 °/s, N=19 flies). We will refer to these induced walking bouts as “opto-runs” to differentiate them from spontaneous walking behavior.

During both opto-runs and spontaneous walking, the presence of visual feedback, selfgenerated visual information, induced walking bouts with on average lower course variability than in darkness (**Suppl. Fig. 1A**). In addition, in darkness, opto-runs displayed on average less course variability than spontaneous walking (**Suppl. Fig. 1A**, compare axis range before and during activation of BPNs). These findings support the hypothesis that the high Vf during opto-runs reflects an (induced) intention to maintain a stable course. Unexpected deviations from this course should therefore recruit activity in premotor steering-control networks, including HS cells. To prevent any visual modulation of HS-cell activity that might confound the activity modulation by Vf, and to prevent vision-based startle responses to the optogenetic light illumination (**Suppl. Fig. 1A**, note the startle response of the fly at onset/offset of the light presentation), we used genetically blinded flies. Like spontaneous walking bouts, HS-cell activity and Vf co-varied at high temporal frequencies (5–10Hz) during opto-runs (**Fig. 2G**). In addition, we found that the correlation between neural activity and Va during opto-runs had the characteristic peak at low frequency and a second peak at a higher frequency range (**Fig. 2G**). We will revisit this specific correlation at a fast timescale later (**Fig. 5**). These observations confirm that the opto-run paradigm is suitable to examine the interaction between HS-cell activity and fine leg movements during high-speed walking bouts.

### HS-cell activity is phase-locked to the walking stride cycle

The fast-timescale correlation between neural activity and Vf suggests that HS cells may be influenced by the stride cycle. To determine the relationship between stride cycle and HS-cell activity directly, we simultaneously tracked the movements of three legs from the left side of the fly ^41^ and recorded the activity of HS cells during opto-runs (**Fig. 3A** and **S1B, C,** see **Methods**). We observed that the oscillatory dynamics of HS cells were strongly coupled to the stride cycle with peak-to-trough amplitude varying from 2 to 6 mV (**Fig. 3B, C** and **S1D,** mean±SEM=4.0±0.3 mV, N=19 fly-cell pairs). Hereafter, to analyze phase relations between neural activity, Vf, and the stride cycle of a leg, we down sampled the data on HS-cell activity and Vf to the leg tracking video’s sampling rate (100 Hz). The oscillations in HS cells were also present in spontaneous, high-speed walking bouts and under visual feedback (**Fig. 3D**). These observations indicate that oscillations may not be induced exclusively by activity of BPNs *per se*, and that they are a property of the dynamics of HS cells even under naturalistic visual conditions.

**Figure 3:**
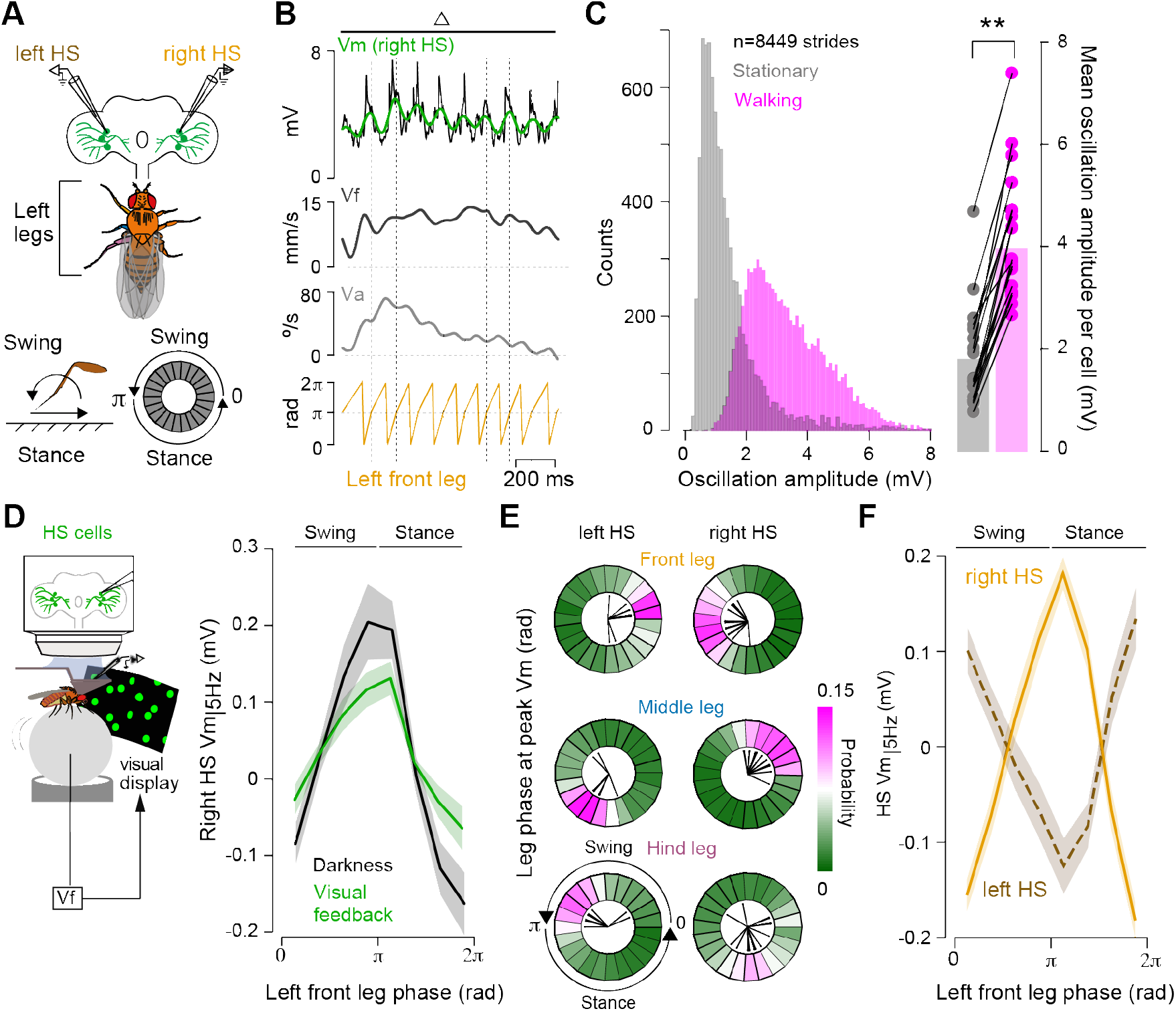
High-speed walking bouts reveal specific phase-relations between neural activity and the stride cycle of different legs. **(A)** Top and middle, definitions for the simultaneous recordings of neural activity and leg movements, color-coded based on leg identity (see also **Suppl. Fig. 1, Video SI).** Bottom left, definition of stride cycle phases. Right, the stride cycle represented in a polar plot, with 0 to π, and π to 2π corresponding to the swing and stance phases, respectively. **(B)** Example segment (triangle indicated in **Fig. 2E)** of simultaneous recordings of neural activity (top, black: raw, green: filtered), the fly’s Vf (black) and Va (gray), and the cycle period of the left front leg (orange), triggered at the transition from the swing to the stance period. (C) Left, distribution of the amplitude of the oscillations during a stride time window (see **Methods)** in walking (magenta) vs. stationary (gray) segments across all fly-cell pairs. Right, mean amplitude of oscillations in walking (3.95±0.29mV) and quiescence (1.79±0.22mV) segments (N=19 fly-cell pairs, walking vs. quiescence: p = 0.000134, z=3.823, Wilcoxon signed-rank test). (D)Left, schematic of the experiment under the presence of visual feedback. Right, tuning of HS cells (high-passed filtered, Vm_|5Hz_) to the left front leg stride cycle during spontanous walking in darkness (black) and under visual feedback (green, N = 8 fly-cell pairs, see STAR Methods). Shaded area indicates SEM. (E) Probability distributions of a leg’s stride cycle phase at the peak of the oscillations of neural activity, displayed on a polar plot. The distributions were calculated by pooling all the strides across all fly-cell pair recordings (contralateral cells, number of strides: 4182–4747 strides, n=19 fly-cell pairs; ipsilateral cells, number of strides: 1937–2269 strides, n=11 fly-cell pairs). Black lines indicate the mean leg phase at peak Vm for each HS cell recording. **(F)** Tuning of HS cells (high-passed filtered, Vm_|5Hz_, right cells: orange solid line, N=19 fly-cell pairs; left cells: maroon dashed line, N=11 fly-cell pairs) to the front leg’s stride cycle during opto runs. Shaded areas indicate SEM.

Under spontaneous or BPN -induced walking conditions, each leg displayed a specific phase relation with the activity of HS cells, which depended on the neural recording side (**Fig. 3D-F** and **S1D–F**). For example, the early stance phase of the left front leg coincided with the peak of the rightside and trough of the left-side HS-cell membrane potential (Vm) (**Fig. 3D-F**). On the spherical treadmill, the left-side front and middle legs exhibited a quasi-opposite relationship, and the corresponding phase relation of each leg with the contralateral, right-side HS cell’s activity was shifted by about 120° (**Fig. 3E**), consistent with a tetrapod-like gait configuration ^38,39,42^. We note that the hind leg’s relationship with neural activity was less robust than that of the other legs (**Fig. 3E** and **S1G–I**), and therefore this was not the focus of further analysis. Altogether, these observations reveal a fixed correlation between a specific leg’s stride cycle and neural activity, and an antiphase relationship between activities in bilateral HS-cell populations.

The oscillatory dynamics of HS cells could also originate from a mechanical coupling between the forces exerted by the legs and movement in the brain of the head-fixed preparation, or as a direct consequence of the activation of BPNs *per se*, independent of walking. To address the possibility of mechanical coupling, we recorded from other genetically identified populations of cells that do not display strong modulation by Vf, such as VS cells ^28^, whose cell bodies are close to those of HS cells. We performed whole-cell patch recordings from VS cells and found that their activity is not coupled to the stride cycle during opto-runs (**Suppl. Fig. 1J–L**). To test the direct effect of BPN activity on HS cells, we decoupled the activity of BPNs from the walking behavior of the fly by stopping the airflow of the spherical treadmill while stimulating BPNs and recording from HS cells. This procedure induced leg movements that were rather uncoordinated. Under these conditions, Vm oscillations were degraded (**Suppl. Fig. 2**), demonstrating that the rhythmic activity of HS cells is driven neither by BPN activation *per se* nor by descending signals responding to BPNs. Thus, the oscillatory activity of HS cells appears to reflect coordinated and periodic leg movements.

### A phenomenological model suggests that the contralateral front leg contributes to the observed oscillations in HS-cell activity

The phase relation between the stride-coupled modulation of HS cells and the fast oscillations of Vf may suggest possible configurations for how leg movements could modulate HS cells. We focused on the movement of front legs since HS-cell excitability (i.e., the temporal derivative of Vm) was more strongly modulated when aligned to the front-than to the middle-leg’s stride cycles (**Suppl. Fig. 3A, B**). Opto-runs with low Va were associated with Vf that displayed two peaks occurring at the stance and swing phases of the stride cycle, reflecting a balanced contribution to acceleration by the left-right pair of front legs (**Suppl. Fig. 3C**) ^38,39,42^. In contrast, when flies deviated from a straight course, the relation between the stride cycle and Vf was single peaked, with the peak occurring at the stance phase of the side dominating the acceleration and driving the direction of angular drift (**Suppl. Fig. 3D**). Regardless of drift direction and amplitude, the peak of the oscillatory dynamics of HS cells always occurred during the same phase of the stride cycle (**Suppl. Fig. 3C, D**). As a result, the peak of the oscillating Vm coincides with the peak of the fast fluctuations of Vf only when the contralateral front leg dominates the fly’s acceleration (**Suppl. Fig. 3D-F**).

Given the phase relation between right-sided HS cells and the stride cycle of the left front leg (**Fig. 3F**), either the left front leg movements could drive the inhibitory phase of neural activity during stance (Model 1, **Suppl. Fig. 4A, left**) or the right front leg movements could drive the excitatory phase of neural activity during stance (Model 2, **Suppl. Fig. 4A, right**). Model 1 predicts that the oscillations of right-sided HS cells and Vf should correlate strongly during leftward drifts because of the shared source for both oscillations (i.e., the left front leg movement) (**Suppl. Fig. 3D** and **S4B,** left), whereas a small and negative correlation is expected during rightward drifts (**Suppl. Fig. 3D** and **S4B,** right). The converse is predicted by Model 2. To examine these predictions, we developed phenomenological models that incorporated the observed phase relations between neural activity, Vf, and front leg movements (**Fig. 3F**, **S3D,** and **S4C, D**, see **Methods**). While either model replicated the observed phase relation between the stride cycle and neural activity by design (**Suppl. Fig. 4E**), the relationship between Vf and Vm in the context of angular drifts was reproduced only with Model 1 (**Suppl. Fig. 4F, G**). The results of these simulations support a model in which a one-sided leg-associated sensorimotor network dominating the acceleration of the fly and thus driving the angular drift configures the opposite-sided HS cells to respond rapidly. The phase analysis of neural activity and Vf, together with the results of the phenomenological models, support the hypothesis that a direct effect of the stride cycle, rather than an indirect effect via Vf, drives the oscillatory dynamics of the population of HS cells.

### Rhythmic activity in HS cells originates from leg-associated sensorimotor circuits

To directly examine the possibility that neural signals from leg sensorimotor circuits underlie the stride cycle-coupled activity in HS cells, we perturbed chemical synaptic transmission via selective expression of the tetanus toxin light chain ^43^ in a large population of leg mechanosensory neurons ^39^ (**Fig. 4A**). Although this manipulation targets chemical but not electrical synapses, spontaneous walking in experimental freely moving flies is typically characterized by curvilinear paths that are not observed under the same behavioral paradigm in control flies, supporting the role of mechanosensory feedback in fine-tuning walking coordination ^28,39^. Recordings from HS cells in experimental flies during spontaneous walking bouts at high Vf (>5mm/s) showed that the oscillatory dynamics coupled to the stride cycle were largely degraded, although not fully abolished (**Fig. 4B, C**). Thus, leg mechanosensory signals are an important component of the oscillatory activity dynamics in HS cells.

**Figure 4:**
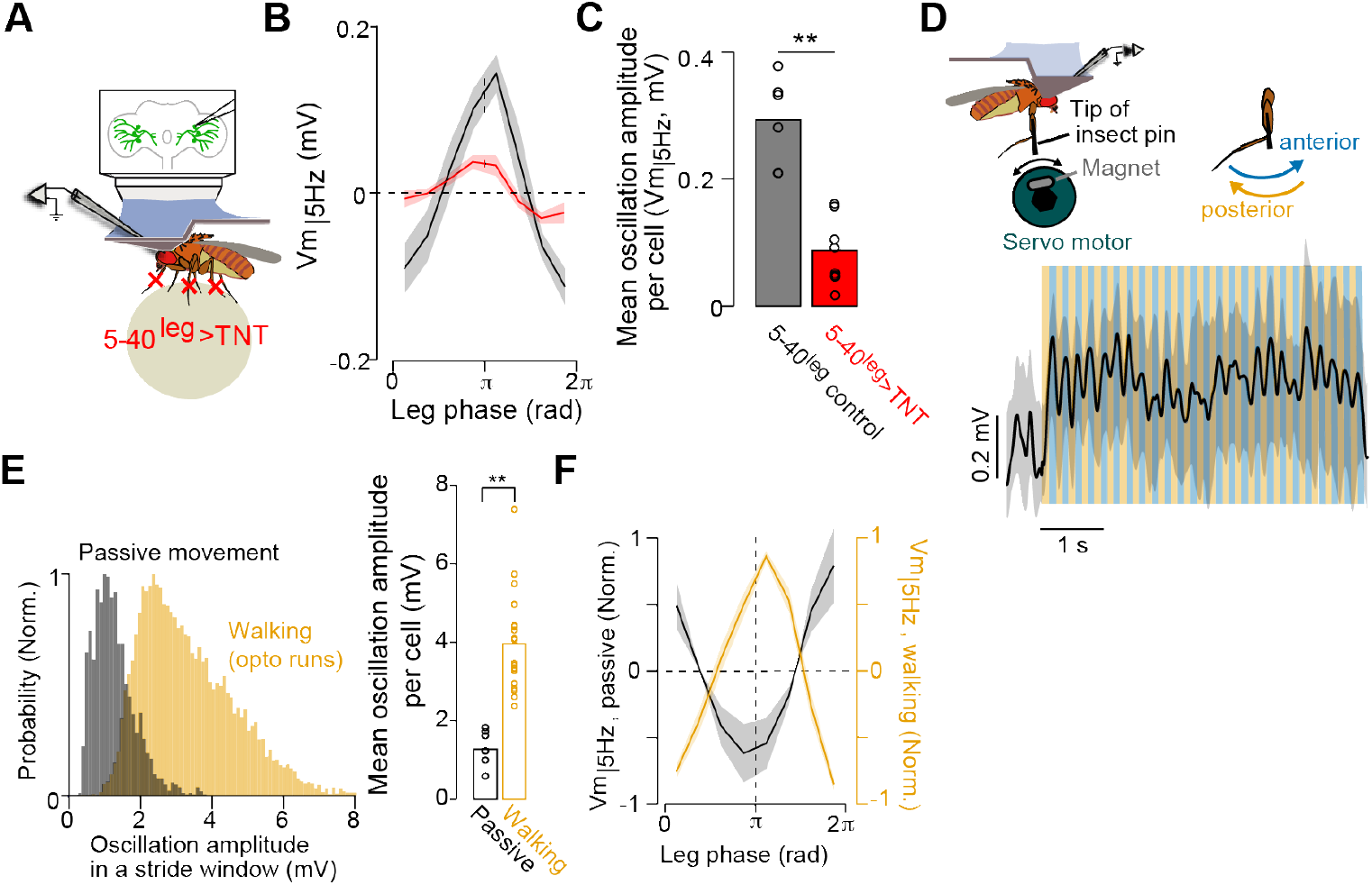
Activity from leg mechanosensory cells contribute to the stride-coupled modulation in HS cells. **(A)** Whole-cell patch recording from right HS cells was performed in spontaneously walking flies with blocked synaptic transmission selectively in leg mechanosensory neurons (“5-40leg>TNT”. see **Methods). (B)** Tuning of right HS cells (high-passed filtered, Vm_|5Hz_) to the stride cycle of the left front leg (grand mean ± SEM). Color code: red, “5-40leg>TNT” flies (N=8 fly-cell pairs); black: control flies (N=7 fly-cell pairs). (C) Averaged magnitude of oscillations calculated as the difference between the peak and trough of the tuning (Vm_|5Hz_) in 5-40leg>TNT vs. control flies (**: p=0.00067, Wilcoxon rank-sum test). (D) Top, schematic of the experimental configuration for passive movement of the left front leg. Bottom, time course of an example experiment. Mean membrane potential (mean Vm ± SD, n=10 trials) during passive oscillations of the leg along the anterior (blue)-posterior (orange) axis. **(E)** Distribution of the amplitude of the oscillations during a stride time window (see STAR Methods) in walking (opto-runs, orange) vs. passive movements of the leg (grey) across all fly-cell pairs (walking: N=19 fly-cell pairs; n=8449 strides; N=8 fly-cell pairs, n=1920 fictive strides). Right, mean amplitude of oscillations in walking (3.95±0.29mV) and during passive movement of the left front leg (1.27±0.15mV) (walking vs. passive movement: p<10^-4^, z=4, Wilcoxon ranksum test). (F) Tuning of the right HS cells to the passive movement of the left front leg (black trace, normalized, Vm_|5Hz_, grand mean ± SEM). For comparison, the orange trace shows the tuning of right HS cells to the stride cycle of the left front leg (normalized Vm_|5Hz_, grand mean ± SEM).

To determine whether leg mechanosensory signals are sufficient to account for the stride-coupled Vm oscillations in HS cells, we passively moved a front leg in a stationary fly via a magnetic system that induced periodic antero-posterior movements of the leg ^44^ (**Fig. 4D**). Although this experimental paradigm does not recapitulate the natural trajectory of the leg during the swing-stance phases of the stride cycle, periodic passive movements of the leg were associated with periodic activity in HS cells (**Fig. 4D**). However, the averaged amplitude of the Vm oscillations was a fraction of that observed during walking (**Fig. 3B, 4E**). In addition, the phase relation between the oscillations of Vm in HS cells and the movement of the contralateral front leg was opposite to that observed during walking (**Fig. 4F**). These differences may arise from the movement of one rather than six legs, the activation of different sensory systems due to the nature and trajectory of the leg movement (for example, no contact with surface), or the lack of additional internal signals that switch the sign of sensory feedback between standing and walking within VNC networks ^18,45,46^. Nevertheless, these findings demonstrate that mechanosensory activity from the legs ascends to circuits in the brain to drive stride-coupled oscillations in HS cells. Additional internal signals are likely combined with the mechanosensory information either within the VNC or brain circuits.

### Stride-coupled modulation tunes neural excitability correlated with rapid steering adjustments

Having established the presence and origin of the stride-coupled oscillations in HS cells, we next asked about their role in course control. One possibility is that they provide a timing signal for rapid steering adjustments. For example, to stabilize the walking direction on a stride-by-stride basis, HS cells and other premotor circuits contributing to course control must regulate their excitability at a precise moment during the stride cycle to be able to generate adjustments in leg parameters that attenuate deviations from a set course direction (**Fig. 5A**). If this were the case, HS cells should display increased activity just a stride cycle before a decrease in angular velocity within the following stride. To test this hypothesis, we examined walking segments displaying deviations from a straight course contraversive to the neural recording side (e.g., left angular drift for right HS cells), since HS cells are excited under this condition ^28^. Recall that during contraversive rotations the peak of the fast fluctuations of Vf coincides with the peak of oscillations in HS cells (**Suppl. Fig. 3F**). This allowed us to examine the activity of HS cells within a time window of 400ms, on average 2 strides during opto-runs (**Fig. 3B** and **S1D**), centered at the peak of the fast Vf fluctuations, when the contralateral front leg dominates the acceleration of the fly (**Suppl. Fig. 3D**).

**Figure 5:**
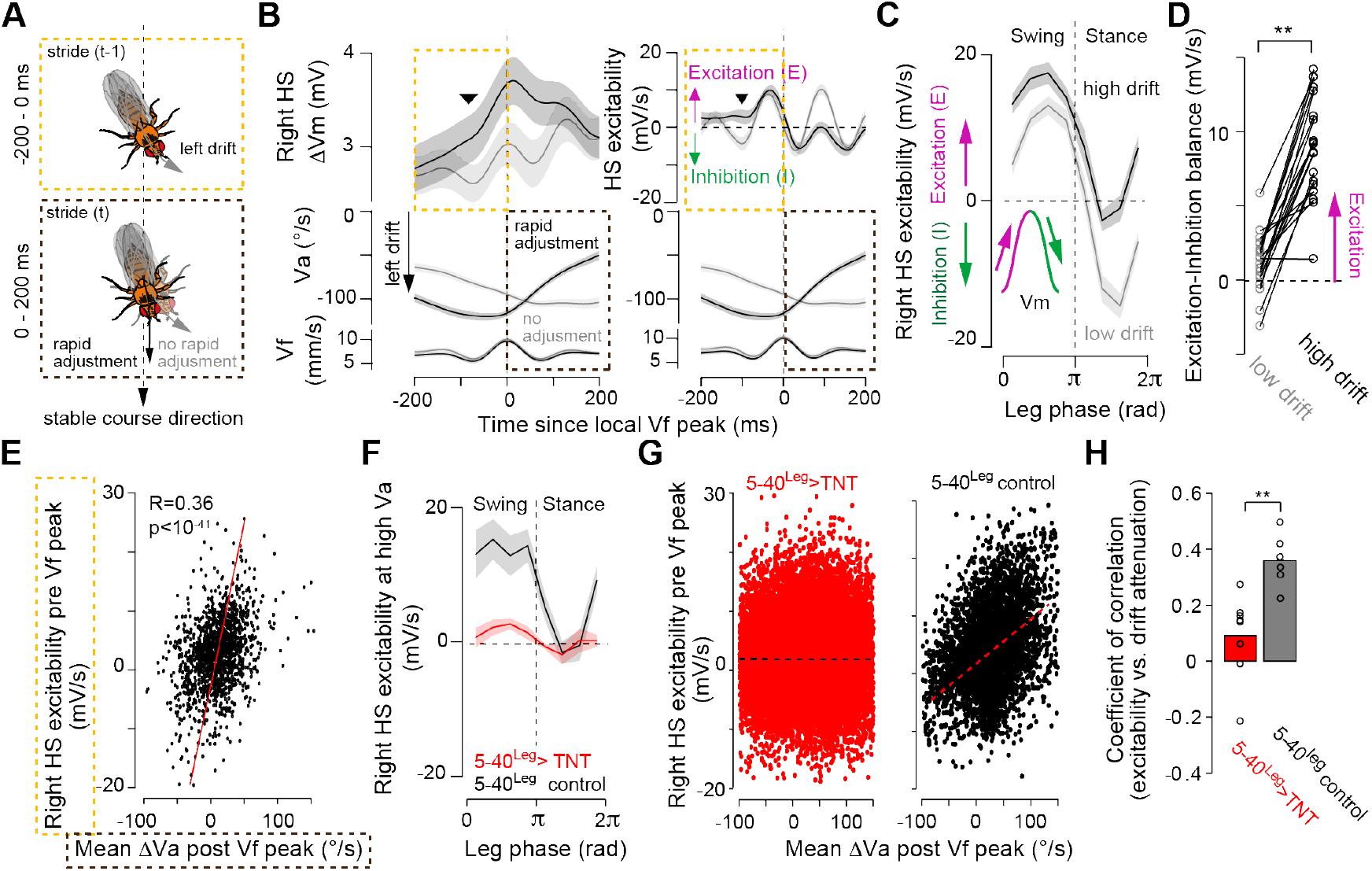
Imbalanced excitation-inhibition in HS cell activity within a stride correlates with steering adjustments in the following stride. **(A)** Schematic with time windows used for analysis. **(B)** The change in Vm in HS cells (relative to quiescence, ΔVm, left), or HS cells’ excitability (right, see **Methods),** and the angular (middle) and forward (bottom) velocities triggered at the local peak of the oscillation of Vf in walking segments with left drift (Va<−50°/s, see A). In both plots, traces were separated based on the magnitude of drift attenuation after the Vf local peak (black or gray, respectively; see **Methods).** Dashed orange and maroon squares indicate the preceding (−200–0ms) and following (0–200ms) time windows from the local Vf peak. Traces and shaded areas represent the grand mean ± SEM, respectively (N=19 fly-cell pairs). The black arrowhead represents a moment in which HS cells were not inhibited within a stride. (C) Tunings of HS cells’ excitability (right-side cells) as a function of the left front leg stride cycle in walking segments with low (gray, −50°/s<Va<~0°/s) or high (black, −200°/s<Va<−150°/s) leftward drift (N=19 fly-cell pairs, left). Positive (magenta) or negative (green) values correspond to excitation or inhibition of the Vm as shown in the inlet. (D) Excitation-inhibition balance across strides in walking segments with low or high angular drift (p=0.00016, Z=−3.78, Wilcoxon signed-rank test, n=19 fly-cell pairs). (E) Mean HS cells’ excitability per walking segment (n=1378 segments from 19 fly-cell pairs) within the preceding time window (−200–0 ms from local peak Vf) as a function of the mean drift attenuation in the following window (0–200ms). The red line indicates the linear regression. (F) Tuning of right HS cells’ excitability during high leftward drift as a function of the left front leg stride cycle (grand mean ± SEM; control, black: N=6 fly-cell pairs, experimental flies, red: N=8 fly-cell pairs). (G) Mean HS cells’ excitability per walking segment (experimental flies, red: n=18319 segments from 8 fly-cell pairs; control flies, black: n=3065 segments from 6 fly-cell pairs) within a preceding time window from the local peak of Vf (−200–0 ms) as a function of the mean drift attenuation in the following window (0–200ms). The dotted lines indicate the linear regression (experimental flies: R=−0.01, p=0.41; control flies: R=0.38, p<10^-16^) (H) Coefficient of correlation between the excitability of HS cells and the magnitude of drift attenuation per fly (experimental flies: N=8 fly-cell pairs; control flies, N=6 fly-cell pairs, **: p<0.0013, Wilcoxon rank-sum test).

To investigate the temporal relationship between activity in HS cells during one stride and the magnitude of Va in the following stride, we separated the time course of neural activity and Va based on the magnitude of rapid adjustments (i.e, a reduction in Va) within the 200ms following the peak of Vf (**Fig. 5A, B**), and then examined the corresponding activity in HS cells within the 200ms preceding the peak of Vf. On average, HS cells were more depolarized at about 100ms before peak Vf in segments with vs. without rapid adjustments after peak Vf (**Fig. 5B,** left, N=19 fly-cell pairs). One likely possibility underlying this effect was a reduction in inhibition as determined by the excitability of HS cells (**Fig. 5B,** right). The increase in excitability for segments with rapid adjustments in Va (**Fig. 5B**) suggests that the excitation/inhibition balance of the neuron within a stride cycle may be rapidly tuned according to the fly’s specific walking state. To examine this possibility, we analyzed the excitability of right-sided HS cells triggered by the left front leg’s stride cycle contingent on the ongoing Va. For walking segments with very low Va (|V_a_|< 50°/s), HS cells were excited during the swing phase and inhibited during the stance phase (**Fig. 5C**), with excitation-inhibition phases that were on average balanced (**Fig. 5D**). In contrast, during walking segments with higher Va(−200 °/s<Va<−150°/s), the excitation phase was longer and the inhibition phase shorter over the stride cycle, resulting in an overall shift to excitation (**Fig. 5C, D**). The effect of the interaction between the magnitude of Va and the stride-coupled modulations, leading to a non-linear amplification in the activity of HS cells (**Fig. 5B**), was largely based on a decrease in inhibitory drive during the stance phase (**Suppl. Fig. 5F**). These findings show that excitation/inhibition balance in HS-cell activity induced by the stride-coupled modulation is fine-tuned by the ongoing magnitude of Va. That is, a component of the state of the body defined by Va interacts with fast, stride-coupled modulations likely to tune activity of HS cells for rapid steering adjustments.

If the stride-coupled tuning of excitability conditioned by the state of Va is important for rapid steering adjustments, then we should observe a correlation between the excitability of HS cells preceding peak Vf and the magnitude of rapid adjustments in Va post peak Vf on a segment-by-segment basis. Indeed, the mean excitability of HS cells preceding the peak of Vf was strongly correlated with the mean decrease in Va post peak Vf on a segment-by-segment basis (**Fig. 5E**, n= 1378 segments, N=19 fly-cell pairs). Importantly, this correlation was independent of the magnitude of Va preceding the peak of Vf (**Suppl. Fig. 5A-E**), which is known to modulate activity of HS cells at longer timescales ^28^. These findings suggest that perturbations in the stride-coupled modulation of HS-cell excitability should degrade the event-based correlation between HS-cell activity and rapid steering adjustments even under conditions with a large Va. We tested this possibility by analyzing the correlation between HS-cell excitability and rapid adjustments on an event-by-event basis in flies with compromised synaptic transmission in a large population of leg mechanosensory neurons that contribute to the stride-coupled modulation of HS cells (**Fig. 4B, C**). Importantly, this manipulation does not affect the modulation of HS-cell activity by Va at longer timescales ^28^. In this manner, we can uncouple the function of stride-coupled modulation from the effects of Va on the activity of HS cells.

Consistent with our previous observations (**Fig. 4B, C**), the excitability of HS cells in experimental flies was very low over the stride cycle, even in walking segments with high ongoing Va (**Fig. 5F**). Furthermore, in experimental flies, we found that the overall excitability of HS cells before peak Vf does not correlate any longer with rapid adjustments post peak Vf (**Fig. 5G,H**). That is, the activity of HS cells is uncoupled from rapid steering adjustments. The remaining adjustments observed in experimental flies implies additional mechanisms that might be explained by the elastic mechanical property of the leg muscles and tendons ^47^ or by other neural mechanisms independent of the activity of HS cells and associated networks. Altogether, these findings demonstrate that stride-coupled modulations provide a timing signal to rapidly change the excitability of HS cells for rapid steering adjustments contingent on the state of Va and that of Vf (**Fig. 1**). We will return later (**Fig. 7**) to the relationship between stride-coupled modulation and the state of Vf. The rapid, fine tuning of HS-cell excitability is controlled by the contralateral leg dominating the acceleration of the fly, thereby contributing to the deviations from a stable course direction at rapid timescales.

### LAL-PS-ANs_contra_ contribute to stride-coupled modulation of HS cells

Based on the contralateral connectivity suggested in model simulations (**Suppl. Fig. 4**) and the consistent functional observations (**Fig. 3, 5**), we sought to identify candidate ANs that could contribute to the timing signals instructing activity of HS cells for rapid steering adjustments. HS-cell axons terminate within the premotor PS ^24^, which is richly innervated by ANs ^26^. Based on the assumption that HS cells receive ascending inputs at the PS (i.e., at their axon terminals), we searched for candidate ANs innervating the PS using an electron-microscopy (EM) hemibrain dataset ^25^. We then applied computational tools to identify the EM-traced skeleton in collections of light-microscopy images from transgenic fly lines ^48^. Following this systematic approach, we identified a pair of previously undescribed ANs that innervate the front and middle leg neuropil, cross the commissure in the VNC, and project to the contralateral PS and a higher premotor region known as the lateral accessory lobe (LAL) (**Fig. 6A** and **S6A–D**) ^24^. Given this projection pattern, we defined this pair of genetically identified ANs as “LAL-PS-AN_contra_”. Even though the axons of HS cells overlap with the axons of LAL-PS-AN_contra_, the pair of ANs do not provide direct chemical synapses onto HS cells (**Suppl. Fig. 6D, E**). Analysis of connectivity indicated that the putative left LAL-PS-AN_contra_ is one to three hops away from each right HS cell (**Suppl. Fig. 6E**). Nevertheless, optogenetic activation of LAL-PS-AN_contra_ induced a robust response in HS cells (**Fig. 6B**), demonstrating functional connectivity. Silencing the activity of these ANs did not affect the fly’s overall walking activity (experimental N=13 vs. control N=12 flies; absolute angular velocity 53.0±2.7 °/s vs. 46.2±3.4 °/s, p=0.11, Z=1.60; forward velocity 3.0±0.3 mm/s vs. 3.3±0.3 mm/s, p=0.43, Z=−0.79, mean±SEM, Wilcoxon rank-sum test). However, silencing the activity of only these two cells was sufficient to induce a significant decrease in the magnitude of the stride-coupled Vm oscillations (**Fig. 6C, D**). These findings support the contribution of LAL-PS-ANs_contra_ to the stride-coupled signal in the functional network of HS cells.

**Figure 6:**
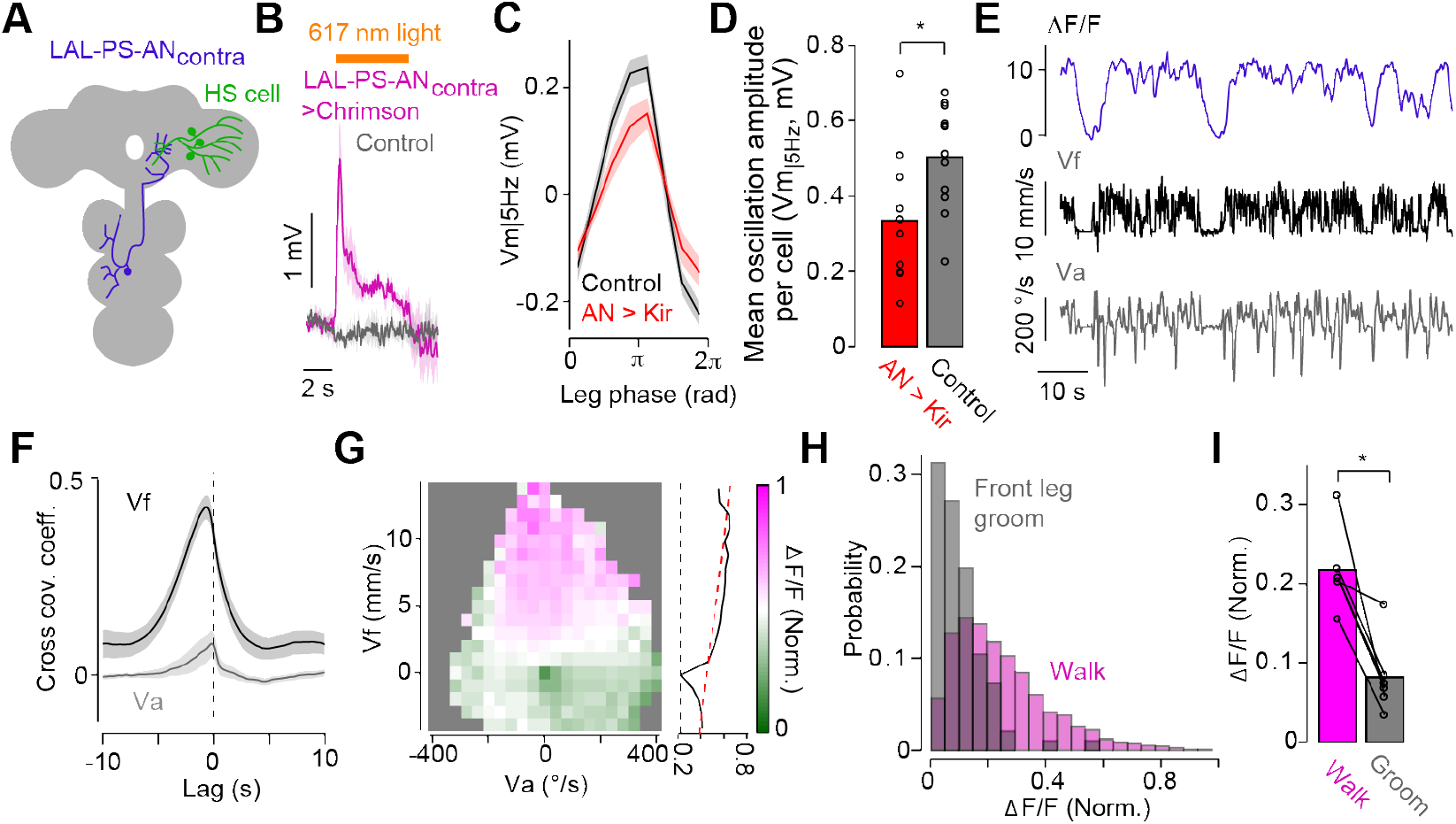
A single pair of speed-sensitive ascending neurons contribute to the stride-coupled modulation in HS cells. **(A)** Schematic of the identified ascending neuron class (violet, LAL-PS-ANs_contra_) projecting close to the axons of HS cells (green) within the inferior posterior slope (IPS). (B) Time course of activity of right HS cells (Vm) in response to activation of LAL-PS-ANs_contra_ via the selective expression of csChrimson (magenta, N=5 flies). Control flies lack expression of csChrimson in LAL-PS-ANs_contra_ (gray, N=3 flies). The orange bar indicates the stimulus duration. Shaded areas indicate SEM. (C) Tuning of HS cells (high-pass filtered, Vm_|5Hz_) to the left front leg stride cycle in flies with LAL-PS-AN_contra_ expressing selectively the inward rectifier channel Kir (red, N=11 fly-cell pairs) and control flies (black, N=12 fly-cell pairs). Shaded areas indicate SEM. (D) Oscillation amplitude of Vm (high-passed filtered, Vm_|5Hz_) calculated as the difference between the peak and trough of the tuning in experimental (left, red, n=11 fly-cell pairs) and control (right, black, n=12 fly-cell pairs) flies (p=0.029, Z=−2.18, Wilcoxon rank-sum test). **(E)** Example time series of relative calcium fluorescence (ΔF/F) of LAL-PS-ANs_contra_ (top), the fly’s forward (middle, Vf) and angular (bottom, Va) velocities. (F) Cross covariance coefficient between left LAL-PS-AN_contra_ calcium signals (ΔF/F) vs. Vf (black) or Va (gray) (N=7 flies). Curves represent the grand mean ± SEM. (G) The activity of left LAL-PS-ANs_contra_ (ΔF/F) as a function of the fly’s Va and Vf. Calcium signals were normalized by the maximum value per fly (N=7 flies). On the right of the map, the Vf tuning (mean calcium signals over Va) was plotted with a linear fitting (the dashed red line). (H) Probability distributions of LAL-PS-AN_contra_ activity (ΔF/F) during walking (magenta, 2909 events) vs. front leg grooming (gray, 96 events). Data were collected from 6 flies. (I) Mean calcium response (calculated per fly) during walking vs. front leg grooming (N=6 fly-cell pairs, p=0.031, Wilcoxon signed-rank test).

To evaluate the response properties of LAL-PS-AN_contra_, we performed 2-photon imaging of the internal free calcium dynamics at their axon terminals in walking flies (**Fig. 6E**). Activity of LAL-PS-AN_contra_ was positively correlated with Vf and relatively insensitive to Va (**Fig. 6E–G**). Moreover, the activity of these ANs was significantly greater during walking than during grooming (**Fig. 6H, I**). Altogether, these findings reveal a pathway that informs about coordinated movement representing Vf and not Va, and that connects stride-coupled signals from the VNC to oscillatory activity in visuomotor circuits in the brain.

### The stance duration over several strides provides an internal motor context

The stride-coupled modulation of HS cells provides a timing signal to rapidly change their excitability for steering adjustments contingent on the state of Va (**Fig. 5**) and that of Vf (**Fig. 1**). Because the activity of LAL-PS-AN_contra_ is correlated with the fly’s Vf and not Va, and silencing their activity decreased the stride-coupled modulation, stride-coupled modulation could also contribute to the representation of the state of Vf at timescales longer than a stride. Our data are consistent with the hypothesis that the stance phase of the contralateral front leg provides an inhibitory drive to HS cells (**Fig. 3, 5** and **S4**). Stance duration is known to be inversely correlated with Vf in freely walking animals ^38,39,49^, as well as in our head-fixed flies walking on a ball (**Fig. 7A, Suppl. Fig. 7A**). Therefore, we reasoned that the duration of the stance over sequential strides could drive an accumulating modulatory signal representing the state of Vf, the motor program linked to the behavioral goals of the fly, or both. That is, the duration of the stance phase integrated over several strides would create a motor context to flexibly recruit the activity of HS cells for rapid steering adjustments. Consistent with this hypothesis, we found that HS cells were on average more hyperpolarized at the end than the beginning of a cycle of strides with long stance duration, and this effect was not observed in strides with short stance durations (**Fig. 7B, D**). If the fly executes sequential strides of equal or similar stance duration, then the modulatory effect per stride would increase to promote a slow overall hyperpolarization or depolarization in the activity of HS cells. Indeed, we found that stance duration modulates the overall depolarization state of the cells over the course of similar sequential strides (**Fig. 7C, D** and **S7B, C**). Consistent with the proposed underlying connectivity structure (**Suppl. Fig. 4** and **6**), this slow modulation of neural activity was specific when aligned to the contralateral but not the ipsilateral front leg stride cycle (**Suppl. Fig. 7D–G**).

**Figure 7:**
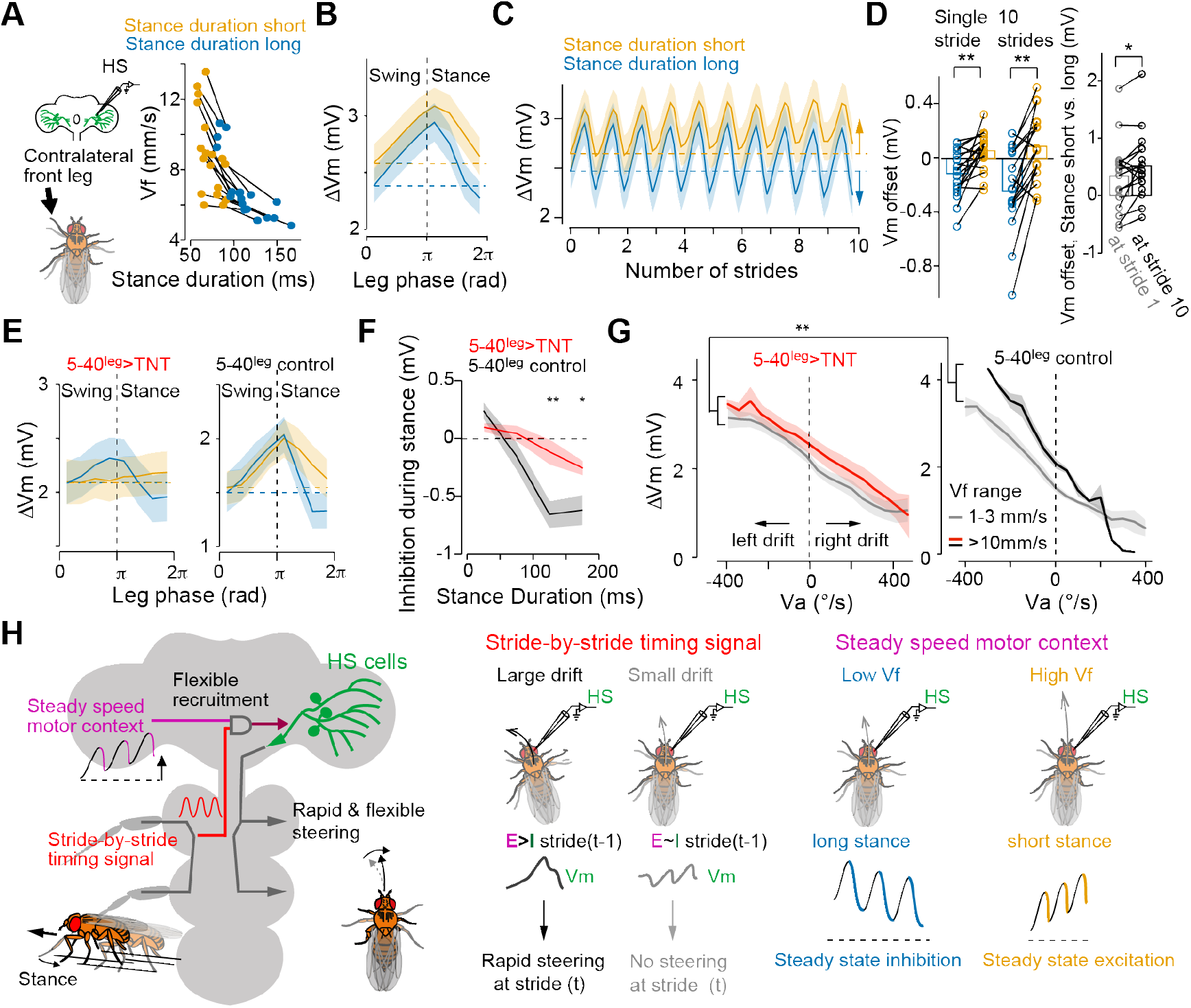
Stance duration controls the inhibitory modulation in HS cells at multiple time scales. **(A)** Left, schematic of the experimental configuration. Right, the relationship between Vf and stance duration. Orange: short-duration stances. Blue: long-duration stances (see **Methods** for definition, N=19 fly-cell pairs; data from the same individual relate to a line). Vf was significantly higher when the stance duration was shorter (p=0.000163, Z=3.82, Wilcoxon signed-rank test). **(B)** Tuning of HS cells (ΔVm, relative to quiescence) to the front leg stride cycle for shorter (orange) and longer (blue) stance durations (grand mean± SEM, N=19 fly-cell pairs). (C) Same as in (B), but for ten similar continuous strides (N=17 fly-cell pairs). (D) Vm offsets at the end of a single stride (left), or ten consecutive strides (middle) relative to the beginning (dashed lines in **(B)** and **(C)).** p=0.00054, Z=−3.46, N=19 fly-cell pairs for a single stride; p=0.00060, Z=−3.43, N=17 fly-cell pairs for ten strides, Wilcoxon signed-rank test. Right, Vm offset difference between walking segments with short vs. long stance durations at the end of stride 1 (gray) or stride lO(black). p=0.019,Z=−2.34,N=17 fly-cell pairs, Wilcoxon signed-rank test. **(E)** Same as in **(B)** but in flies with blocked chemical synapses in leg mechanosensory neurons (left, N=8 fly-cell pairs) and the corresponding genetic controls (right, N=6 fly-cell pairs). (F) Inhibition during stance as a function of stance duration (experimental flies, red, N=8 fly-cell pairs; control, N=6 fly-cell pairs). *: p<0.05; **: p<0.01, Wilcoxon rank-sum test. Curves represent the grand mean ± SEM. (G) ΔVm of HS cells as a function of Va during low and high-speed walking in experimental (left, N=8 fly-cell pairs) and control (right, N=5 fly-cell pairs) flies. Vf-dependent Vm offset in experimental flies was significantly smaller than in control flies (**: p=0, bootstrapping method, see **Methods).** Curves represent the grand mean ± SEM. (H) Summary of the findings of this study.

If the emergence of the motor context in HS cells originates from the same source as the rapid tuning of cell excitability (**Fig. 5**), then perturbing chemical synapses in leg mechanosensory neurons should also disrupt this longer-timescale neural modulation. Evaluating the activity of HS cells in experimental flies showed that the stance-related inhibition was significantly decreased (**Fig. 7E, F**). Consistent with the idea that the stance-based inhibitory mechanism configures motor-context signals in HS cells (**Fig. 1B**), the forward speed-dependent depolarization in HS-cell activity was significantly decreased (**Fig. 7G, see also Fig. 1**). Altogether, these findings demonstrate that the stance duration over sequential strides configures a motor context signal reflecting the state of Vf and of motor programs at play over longer timescales than a stride, thereby explaining how HS cells can be flexibly recruited to contribute to steering control.

## Discussion

A critical component of adaptive behavior is the provision of continuous feedback on the relationship between behavioral goals and the state of the body, which defines a dynamic internal motor context, and the environment. The questions of how such motor contexts emerge and support flexible sensorimotor mapping at multiple timescales during online movement corrections has remained poorly understood. By recording neural activity at high temporal resolution alongside analysis of leg movements, we have demonstrated that HS-cell activity is modulated by stride-coupled signals at multiple timescales to flexibly recruit their activity for rapid steering maneuvers. Over a stride, the rhythmic modulation sets short timing windows upon which the excitation/inhibition balance in the cells’ activity is fine-tuned contingent on the state of Va for rapid course adjustments. At a longer timescale, over several strides with similar stance duration, the modulation builds up, reflecting a motor context that represents a quasi-steady state of Vf. Altogether, our findings establish that stride information from ascending pathways configure multi-timescale contextual signals during walking to flexibly recruit activity in visual circuits for rapid course correction in the appropriate motor program.

### Rapid stride-coupled modulation in the central brain

Our finding of stride-related signals in a class of *Drosophila* visual neurons, the HS cells, provides important insights into the source and function of such rapid sensorimotor signals in brain circuits, and more generally, into the mechanisms of motor context modulation in brain circuits via the critical role of ANs from the VNC. Over the last decade, a considerable amount of work in rodents and insects has revealed prominent modulations in the activity of visual circuits during walking ^50–54^. However, no previous work has addressed the timescales at which these modulations operate. Our study reveals the presence of short-term, stride-coupled signals in visual circuits that are further modulated by walking speed over longer timescales. Thus, in the context of online course corrections, stride-related signals operate at multiple timescales.

In the mammalian cerebellum and motor cortex, although the presence of locomotion-related stride-coupled dynamics is well known, their functional contribution to the control of locomotion is less clear ^21^. Therefore, the computational role of the plethora of extra-retinal, locomotor-related signals in brain sensorimotor circuits has remained an open question. We propose that the anatomical (**Fig. 6** and **S4**) and physiological (**Fig. 3** and **5**) properties of stride-coupled modulation in *Drosophila* HS cells are well conFig.d for a stride-by-stride course correction with proper timing and direction selectivity. First, during its stance phase, a leg contributes to a torque relative to the center of body mass that may induce unintentional body deviations in a direction ipsilateral to the leg ^55,56^. Stride-coupled modulation from a contralateral leg provides a timing signal to HS cells and other elements of the network to correct for such a drift in the next stride (**Fig. 5**), perhaps by modulating the placement of the leg ipsilateral to the neuron during its swing phase ^56^. This contralateral wiring is reminiscent of the lamprey postural control system, where vestibular activations upon yaw swimming drift drive contralateral reticulospinal neurons to trigger corrective steering ^57^. Second, the excitation/inhibition balance in the rhythmic stride-coupled modulation is fine tuned to amplify cell activity contingent on the ongoing state of Va, leading to on-demand, scalable depolarization within a stride cycle (**Fig. 5** and **7H**). With these specific anatomical and physiological configurations, stride-coupled signals in HS cells can directly contribute to the rapid course correction with proper timing and direction selectivity on a stride-by-stride basis (**Fig. 5** and **7H**). Therefore, our data not only reveal the presence of ascending stride-coupled signals in a brain visuomotor circuit but also their functional role directly linked to online course adjustments.

### Neural representation of a motor context

Our results show that the same source that drives stride-coupled modulation in visual neurons configures a slower modulation emerging over several strides (**Fig. 7H**). Long-lasting internal states specified by experience, physiology, arousal and emotions influence patterns of neural activity to coordinate behaviors adaptively according to the needs of an animal ^9,12,14,50,51,58–62^. Internal states are also defined by motor context, the continuous feedback on the relation between behavioral goals and the state of the body. During locomotion, motor-related signals are broadcast throughout the brain, creating “motor contexts” upon which neural activity is tuned in accordance with the perceptual and motor requirements of the animal moving through space ^7,28,63–70^. While internal state-dependent modulations in neural activity are typically based on global gating mechanisms characterized by state transitions, such as behaving vs. resting ^50,51,58,71^, the effect of locomotor-specific signals on neural activity is graded, continuous, and co-varying with locomotor speeds and modes ^28,65,67,68,72,73^. Neural representations of forward speed, a motor context that can be linked to an animal’s vigor to achieve a goal ^74^, have been widely found in rodents ^53,72,75–79^, insects ^28,80^, worms ^65,81^, and fish ^82^. The origin of these speed-related signals, however, remains understudied. Our data describe the source of a forward speed representation in a *Drosophila* visual circuit that ascends from the VNC and explains the mechanism underlying its function based on stance-related inhibition (**Fig. 7**). Because stance duration is inversely correlated with forward speed, continuous, stance-dependent inhibitory modulation builds a representation of the fly’s forward speed and/or behavioral goal in the cells’ activity (**Fig. 7H**). Therefore, our work reveals a mechanistic link between a movement parameter regulating walking speed (i.e., stance duration), a speed representation in the central brain, and their effect on neural activity and steering control.

## Outlook

Our data on the flexible recruitment of a visuomotor circuit for course control based on motor context will be widely applicable to neural circuits controlling behavior during active sensation, navigation, foraging and other tasks requiring flexible mapping between sensory information and movement parameters. Speed-related signals such as the stride-coupled modulation described here are critical for vector computation in path integration ^77,83^ and egocentric-allocentric coordinate transformations ^78,84^. A mechanism integrating the stride-coupled activity over several strides (**Fig. 7**) could be applied as a step counter for path integration, whose neural substrate should exist in the insect brain ^78,84–86^. Therefore, we speculate that similar stride-coupled signals may be transmitted to higher-order brain circuits involved in navigation and motor control ^87–89^. This hypothesis is supported by the finding that LAL-PS-AN_contra_ innervate leg neuropils in the VNC and project to the PS and LAL (**Fig. 6** and **S6**), two major insect brain areas implicated in motor control. The LAL is known as the hub connecting navigational and motor circuits ^24,90,91^. Interestingly, studies in insect olfaction suggest that periodic oscillatory neural dynamics are useful for memory association ^92,93^. The frequency range of the stride-coupled modulation described here is within the theta rhythm (4–12 Hz) (**Fig. 2** and **S1**), where the oscillation could optimally promote spike timing-dependent plasticity ^94^. Future research will reveal how our findings can be generalized and further extended to other circuits in flies and corresponding systems in other species for motor control, navigation, and learning.

## Acknowledgements

We thank Nélia Varela for stock construction and maintenance, and immunostaining; Sebastián Malagon for the analysis with DeepEthogram, and Ibrahim Taştekin, Gregory Jefferis, and Alexander Bates for technical support for image registration. We are grateful for helpful discussions and comments on the manuscript by Maximilian Jösch and members of the Chiappe laboratory. We thank the labs of S. Bidaye, G. Card, M. Dickinson, C. Mendes, V. Ruta, and M. Silies for fly stocks. This work was supported by the Champalimaud Foundation and the research infrastructure Congento, LISBOA-01-0145-FEDER-022170, co-financed by Fundação para a Ciência e Tecnologia (Portugal) and Lisboa2020, under the PORTUGAL 2020 Agreement (European Regional Development Fund). This work was also supported by the Japanese Society for the Promotion of Science (JSPS) Overseas Research Fellowship 20170687 (T.F.), by the Bial Foundation grant 191/12 (M.E.Ch), by the Marie Curie Career Integration Grant PCIG13-GA-2013-618854 (M.E.Ch), and by the European Research Council ting Grant ERC-2017-STG-759782 (M.E.Ch).

## Author Contributions

Conceptualization, T.F. and M.E.Ch; Methodology, T.F.; Investigation, T.F., and M.B.; Formal Analysis, T.F., M.B., and M.E.Ch., Supervision M.E.Ch.; Writing, T.F., and M.E.Ch.

## Methods

### Fly husbandry

Flies (*Drosophila melanogaster*) were reared in standard medium at 25°C with a 12-hr light and 12-hr dark cycle. We noticed that male flies have more robust walking behavior on the ball than females. For this reason, we randomly selected male flies for all experiments. We excluded flies that looked unhealthy at the time of the fly preparation, or that those that display less than 10 walking bouts. Overall, this represented about 35% of flies that were discarded for spontaneous walking experiments, and 30% under opto-runs. For the leg mechanosensory perturbation experiments, because of the likely function of these sensory neurons in maintaining coordinated locomotion, many flies walked poorly and never reached the high-speed walking (>5 mm/s for at least 30s, 0.12 g weight of the ball), and we needed to exclude about 70% of flies. For the remaining 30% of flies with this genotype, individual flies displayed walking bouts at high speed that conformed with the threshold criteria. For optogenetic experiments, flies (including controls) were fed with 1 mM all-trans-retinal after eclosion and were kept subsequently in darkness until the experiment. All the experiments were performed with 1-to-4-day-old male flies. Specific sources of transgenic lines are listed in the Key Resources Table.

### Electrophysiological and calcium recordings

Details of the fly preparation for simultaneous physiology and behavior, and the treadmill system are described in ^28,95^. Briefly, a cold-anesthetized fly was mounted on a custom-made holder, and the back of the head’s cuticle was removed with fine tweezers. The dissected fly was mounted under the microscope and positioned on an air-suspended 9 mm diameter ball. *In vivo*, whole-cell patch-clamp recordings and calcium imaging were performed using an upright microscope (Movable Objective Microscope, Sutter) with a 40× water-immersion objective lens (CFI Apo 40XW NIR, Nikon). The external solution, which perfused the preparation constantly, contained 103 mM NaCl, 3 mM KCl, 5 mM TES, 8mM trehalose, 10 mM glucose, 26 mM NaHCO_3_, 1 mM NaH_2_PO_4_, 4 mM MgCl_2_ and 1.5 mM CaCl_2_ (pH 7.3 when equilibrated with 95% O_2_/ 5% CO_2_; 270–280 mOsm). Patch pipettes (5–7 MΩ) were filled with an internal solution containing 125 mM aspartic acid, 10 mM HEPES, 1 mM EGTA, 1 mM KCl, 4 mM MgATP, 0.5 mM Na_3_GTP, 20 μM Alexa 568–hydrazide-Na and 13 mM biocytin hydrazide (pH 7.3; 260–265 mOsm). The neural lamella was ruptured by the local application of collagenase IV ^59^. Current-clamp data were filtered at 4 kHz, digitized at 10 kHz using a MultiClamp700B amplifier (Molecular Devices), and acquired with Ephus. The recorded cell membrane potential (Vm) was corrected for junction potential (11 mV). For HS cell recordings performed in darkness during spontaneous walking (19/25 cells, **Fig. 1B, 2B–D, S3F,** and **S4G**) and for HS cells recordings performed under leg mechanosensory neurons’ perturbations, (**Fig. 4B, C, 5F– H,** and **7E–G**), data was obtained during a previous study (Fujiwara et al., 2017). Synaptic activity from leg mechanosensory neurons was inactivated by the selective expression of UAS-FRT-stop-FRT-TNT in the leg imaginal disc (dac^RE^-FLP, ^96^), driven by 5-40-GAL4, a pan sensory neuron driver ^97^. The rest of the data was collected during this study.

We recorded internal calcium dynamics from the axons of LAL-PS-AN_contra_ projecting at the PS, the neuron’s most superficial projection field, using a custom-built 2-photon laser scanning system. We used a Chameleon Ultra II Ti-Sapphire femtosecond laser (Coherent) tuned to 930 nm for GCaMP excitation (6 mW under the objective lens). Emission was collected on GaAsP PMT detectors (Hamamatsu, H10770PA-40) through a 535/50 nm bandpass filter (Chroma). A 128×128 pixels slice image was acquired at 15 Hz with ScanImage.

### Chemogenetic silencing of HS cells

An electrode filled with external ringer solution containing 1 mM histamine and 40 μM Alexa 568 was placed in juxtaposition to the axon terminal of GFP-tagged HS cells guided by 2-photon imaging. The neural activity manipulation was conditional to the fly’s forward velocity *via* a closed-loop system. Real-time treadmill signals (< 10ms delay) were detected with a panel display controller unit (IO Rodeo, Reiser and Dickinson, 2008). Once the forward velocity reached a threshold (>1 mm/s on average for 3s), brief pulses of histamine (10ms, 6 psi) were applied. Control flies with no artificial expression of ort showed minimal inhibition upon histamine application (**Fig. 1D, F**), indicating that HS cells do not express high levels of ort endogenously. In contrast, brief pulses of histamine induced a reliable hyperpolarization in HS cells expressing ort (−11.9±0.72 mV at the peak, mean±SEM, n=12 cells).

Injecting 1mM histamine at the HS-cell axon terminals also induced a slight but measurable inhibition in VS cells (2.8±0.8 mV n=5 cells, Mean±SEM) that could contribute to the overt effect on steering of the walking fly. To test the contribution of VS cells to behavior, we examined the fly’s behavior when we applied histamine directly onto VS instead of HS axons using the same GAL4 driver. For this purpose, we adjusted the histamine concentration in the solution (300 μM) to induce inhibition in VS cells with a magnitude that was comparable or slightly higher than the one observed when targeting HS axons (−5.8±1.0 mV, n=6 cells, Mean±SEM). Directly inhibiting VS cells in this manner induced no overt effect on the walking behavior of the fly under an identical closed-loop configuration between the fly’s forward speed and the histamine application (**Fig. 1G**). We therefore concluded that despite the off-target expression of the VT058487-GAL4 line, the results from these experiments altogether support the model that the steering effects are a consequence of the unilateral perturbation of the activity of the population of HS cells.

### Optogenetic activation of Bolt protocerebral neurons (BPNs), neural recording, and visual stimulation

Unless otherwise stated, whole-cell patch recordings of HS and VS cells were performed in head-fixed, blind flies (NorpA mutant, ^98^) walking on the spherical treadmill. A fiber-coupled light (617nm, M617, Thorlabs) was projected onto the central part of the brain through the objective lens to activate selectively BPNs expressing UAS-csChrimson. Each trial consisted of 5s of stimulation with light pulsed at 100 Hz (50% duty cycle) and intensity ranging from 42 to 135μW/mm^2^. For **Fig. 6B**, ascending neurons were activated with the fiber-coupled light (36 μW/mm^2^). Walking movements of the left-side front, middle, and hind legs were simultaneously captured with a monochrome digital camera (UI-3240CP-NIR-GL, iDS), coupled to a 25mm focal length lens (M2514-MP2, Computar), and an extender (EX2C, Computar). Images were acquired at 100Hz by externally triggering individual frames from a data acquisition card (USB-6229, National Instruments).

For the subset of recordings labeled as the “Ball stopped” condition (**Suppl. Fig. 2**), the airflow of the spherical treadmill was turned off, thereby making rotations of the ball difficult for the fly. Under this condition, flies stopped locomotion and transitioned into either quiescence or non-locomotive movements. After this “Ball Stopped” condition, the airflow was resumed under the condition labeled as “After”. For analysis, we used six trials of each condition per fly-cell pair (**Suppl. Fig. 2B, C**).

To examine whether the fly’s forward walking induced by BPNs activation was sensitive to course stabilizing visual feedback (**Suppl. Fig. 1A**), we presented the head-fixed fly with a visual stimulus (9° random dots, 16% density) using a 32×96 arrays of 570nm green LEDs (Bet Lux Electronics, Reiser and Dickinson., 2008) in unity gain closed-loop with the fly’s rotation. To activate BPNs non-invasively, we placed an optic fiber (200μm core, M25L02, Thorlabs) 5mm apart from the fly’s head and illuminated it at 100Hz (10% duty cycle) and average intensity of 20μW/mm^2^. Note that the activation strength was turned down to prevent masking the visual stimulus.

To examine whether HS cells activity under the presence of visual feedback was also modulated by stride-related signals (**Fig. 3D**), we recorded HS-cell activity under a unity gain of closed-loop translational visual stimulus (9° random dots, 16% density) coupled to the forward velocity of the spontaneously walking fly. The treadmill signal, sampled at 4 kHz, was integrated over 1.6ms and sent to the LED arena controller to control the translational motion in the visual display (the total closed-loop delay < 10ms). Note that in these experiments, we split the visual display in two, centered at the front of the fly to induce the translational feedback condition. That is, the closed-loop configuration was 1D and did not incorporate the rotations of the fly that may otherwise interfere with the visual responses of the cell, masking the presence of the forward-velocity associated oscillations. To compare the stride-related signals in the same individual and cell under different light conditions, the protocol alternated 2 min recordings between visual feedback and darkness.

### HS-cell recording with external leg movement

To passively move a leg, we adopted the method described in ^44^. Briefly, a tip of a tiny insect pin (12.5μm tip diameter, 26002-10, Fine Science Tools) was inserted into the coxal part of a front leg. A small, rare metal magnet (1×1.5×2 mm, Magnet Solutions) attached to a screw head was placed under the leg and was rotated periodically along the front-back axis of the fly’s body with a servomotor (period: 5Hz, amplitude: 20°). The other legs were removed to avoid the contribution of spontaneous leg movements to HS cells’ activity.

### Immunostaining

Isolated brains were fixed for 30 min at room temperature in 4% paraformaldehyde in PBS, rinsed in PBT (PBS, 0.5% Triton X-100 and 10 mg/ml BSA), and blocked in PBT + 10% NGS for 15min. Brains were incubated in primary antibodies (1:25 mouse nc82 and 1:1000 rabbit antibody to GFP) at 4°C for three days. After several washes in PBT, brains were incubated with secondary antibodies (1:500 goat-anti rabbit: Alexa Fluor 488 and 1:500 goat-anti mouse: Alexa Fluor 633) for three days at 4°C. Brains were mounted in Vectashield, and confocal images were acquired with a Zeiss LSM710 scope with a 40× oil-immersion or 25× multi-immersion objective lens. For the MultiColor FlpOut experiments (**Suppl. Fig. 6**), we followed the original protocol described in ^99^, except 10% NGS was used for blocking steps.

### Data processing and analysis

We used MATLAB (MathWorks, Inc., Natick, MA) for data analysis, neither with specific randomization nor blinding.

### Processing of electrophysiology and behavior data

Electrophysiological and treadmill signals were down sampled to 500Hz and smoothed using a lowess algorithm with a 120ms window. To compare neural and velocity signals to leg movements, time series were further down sampled to 100Hz to match the leg tracking’s video sampling rate. Note that downsampling, filtering, and averaging neural activity degraded the amplitude of neural modulation. Therefore, to examine the amplitude of the modulation (**Fig. 3C, 4E**), we obtained the distribution of the peak-to-trough magnitude of the oscillations within a stride window (from stance onset to the next stance onset) from the original signals. The Vm oscillation within a corresponding short window during stationary periods (i.e., moments with zero treadmill signals before BPNs activation) was calculated to estimate the noise level.

#### Definition of spontaneous walking bouts

In spontaneous walking bouts (**Fig. 1B, 2B–D, S3F, and S4G**), walking-related signals in HS cells were analyzed following ^28^. Briefly, we extracted walking bouts from the treadmill signals using a supervised machine-learning algorithm JAABA ^100^ based on side-view videos of the walking fly. Isolated walking bouts and the corresponding Vm signals were concatenated per fly. For **Fig. 2D, G**, the angular (Va) and forward (Vf) velocities and Vm were bandpass filtered (0.5Hz bin), and the maximum cross-covariance coefficient between the filtered Va and Vm or between the filtered Vf and Vm was calculated at each bandpass range.

#### Bootstrap analysis of modulation of forward velocity on HS cell activity

For statistics in **Fig. 1B**, 1% of all data was randomly chosen (bootstrap) to fit linear regression, and the offset of the fitted line was measured. This procedure was performed both in low vs. high forward speed conditions, and the offset difference was calculated. This was repeated 1000 times to obtain a distribution for the offset difference. The p-value was determined by counting how many data points (out of 1000) crossed the zero value. Similarly, for **Fig. 7G**, the offset difference was calculated for the experimental and control groups, and the p-value was determined by counting how many times (out of 1000 repetitions) the offset value for the experimental was larger than the one for the control. For **Fig. 1E, F**, histamine injection triggered traces were divided depending on the mean value of the forward velocity of the fly within a second before histamine application (−1–0s window).

#### Effect on course control of unilateral, conditional perturbations of neural activity

To quantify the change in the course direction of the fly (**Fig. 1D–G**), we calculated the mean angular velocity within a window of 2s before histamine injection and subtracted it from the mean angular velocity within 2s after histamine injection.

### Effect of the stride-coupled signals on HS cells activity and rapid steering maneuvers

We extracted the excitability of HS cells as the temporal derivative of the Vm (**Fig. 5B**). The change in Vm (relative of quiescence, ΔVm), the excitability of HS cells, and the Va and Vf were triggered at local peaks of the high-pass filtered Vf (>5Hz, Vf_|5 Hz_) during opto-runs. Walking was defined with a threshold of 1mm/s. For each fly-cell pair, these event-triggered segments of data were further classified based on the magnitude of angular drift attenuation, the decrease of Va post peak of Vf. If the magnitude of the decrease in Va was within the 1^st^ or 4^th^ quartiles of the distribution across opto-run segments, these classes were labeled as the low vs. high drift attenuation conditions, respectively. Drift attenuation was defined as the mean Va within a 0–200ms window (peak Vf at 0) subtracted from Va at time 0 (at local Vf peak).

For **Suppl. Fig. 3F**, Vm and Vf were triggered at local peaks/troughs of Vf_|5Hz_ with a threshold of ±1mm/s during left (Va <−50°/s) or right (Va >50°/s) angular drifts. For **Suppl. Fig. 4G**, Vf_|5Hz_ was projected onto a 2D behavioral map (Va bins: 20°/s, Vf_|5Hz_ bins: 0.3mm/s). The value at each pixel was calculated as the mean of all collected data points.

For **Suppl. Fig. 5**, “residual drift attenuation”, drift attenuation that is independent of the mean magnitude of Va within the preceding 200ms window from peak Vf was calculated as follows:

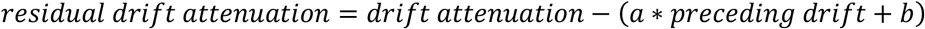

where *a* is the slope coefficient and *b* is the intercept of the linear regression between the drift attenuation and the preceding mean magnitude of Va. For **Suppl. Fig. 5E,** HS excitability, Va, and Vf were triggered at local peaks of Vf_|5Hz_ as described above, and the triggered traces were classified based on the magnitude of the residual drift attenuation following the same class definition as above.

### Leg tracking

We used a machine learning-based strategy (DeepLabCut, ^41^) to monitor leg movements by tracking two prominent joints: the femur-tibia and tibia-tarsus. The training was based on manual annotation of these joints in 4 flies (100 frames each, 0.17–0.26% of the total amount of time per fly). The output of the trained network gave the x and y positions of the joints as a function of time. To define leg phases, time courses were first divided into 1s chunks. Chunks with Vf > 5mm/s and with consistent periodic leg movements (periodicity of leg positions was estimated by their autocorrelation function, **Suppl. Fig. 1G**) were used for further analysis (45% of total chunks used). Similarly, for spontaneous walking datasets (**Fig. 3D, 4B, C, 5F–H, 6C, D, and 7E–G**), walking events with Vf > 5mm/s and with consistent periodic leg movements were selected (20% of chunks used). Local inspection of these chunks with periodic leg movements indicated that the stance onset of the legs corresponded to the local minima (i.e., the forward most position) of the horizontal (x-axis), whereas the swing onset corresponded to the local maxima of the horizontal axis (**Suppl. Fig. 1C**). Therefore, the stance and swing phases were defined when the x-axis position of the leg shifted from the local minima to the local maxima and vice versa, respectively. Combining the positional information of the femur-tibia and tibia-tarsus joints (by the square root sum of squares of their horizontal positions, **Suppl. Fig. 1C**) increased the fidelity of the detection of the local maxima and minima in the trajectory. Next, phase values were linearly assigned to the swing (0 to π) and stance (π to 2π) periods. Similarly, local maxima and minima of HS cells’ Vm were detected, and phase values were assigned to the time series so that the local maxima and minima correspond to 0 and π, respectively. The phase relationship between leg movements and HS cells’ Vm (**Fig. 3E**) was characterized by calculating the mean value of the Vm phase subtracted from the leg phase per stride, and the distributions across strides were plotted. For tuning plots, the mean value of the high-pass filtered Vm (Vm_|5Hz_) (**Fig. 3D, F, 4B, F, 6C, 7B, C, E, S1F, L, S3C, D,** and **S7D, F**), HS excitability (**Fig. 5C, F,** and **S5F**) or the Vf_|5Hz_ (**Suppl. Fig. 3C, D** and **S7B**) at each leg phase bin of π/4 was calculated. For **Fig. 7B, C, E** and **S**7**B, D, F**, data in each recording was divided per stride depending on whether the stance duration was shorter (1^st^ quartile of the length distribution) or longer (4^th^ quartile). Tuning plots were generated with these classified data. For **Fig. 7C** and **S7B, F**, the mean stance duration over 10 consecutive strides was used for the classification. For **Suppl. Fig. 3A**, HS-cell excitability was projected onto a 2D leg phase space with a bin of π/4. The value at each pixel was calculated as the mean across all collected data points.

### Simulation

The model has as inputs the leg’s stride cycle phases (see below) and Va (**Suppl. Fig. 4**). The outputs of the model are HS-cells’ Vm and the fly’s Vf (**Suppl. Fig. 4C, D**). For simplification, the model considered the pair of front legs only. A realistic sequence of swing and stance phases for the left and right legs was constructed as follows. A random sequence of stance durations for one leg was initially generated. Following the distribution of swing and stance durations during BPNs activation, which ranged between 30–60ms and 50–150ms, a sequence of swing phases was added with a duration that was proportional to the previous stance duration within a range. Once the swing and stance phases of one leg were defined, the phases for the other leg were generated such that each swing phase was placed within the other leg’s stance period ^38,39,49^.

Next, for the contralateral leg modulation model configuration (Model 1, **Suppl. Fig. 4A–F**), Vm was based on the contralateral leg phase such that the fictive HS cell was excited from swing onset (*θ_min_* = 0) to early stance phase 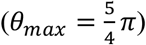 and was inhibited from the early stance phase to the swing onset:

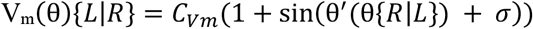

where V_m_(θ){*L*|*R*} is a non-negative value of the left or the right HS cell’s V_m_, θ{*R*|*L*} is the right or the left leg phase, θ′ is an oscillator with values – 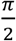 at *θ_min_* and 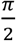 at *θ_max_, C_Vm_* is a constant scaling factor to match the scale of modeled traces to the actual data, and *σ* is a noise parameter.

For the ipsilateral leg modulation model configuration (Model 2, **Suppl. Fig. 4A, C, E, F**), Vm(θ){*L*|*R*} was calculated with the ipsilateral leg phase θ{*L*|*R*} and θ′ became 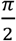 at *θ_min_* and – 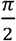 at *θ_max_* (i.e., assumption of excitation during stance).

Based on the observed Vf tuning to the leg phase (**Suppl. Fig. 3C, D**), both the left and right leg phases contributed to Vf in the model, and their specific contribution as a function of the stride cycle was weighted depending on Va. The contribution function of each leg to Vf, F(θ){*L*|*R*} was defined as:

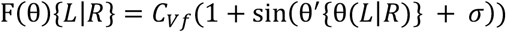

where *θ* is the leg phase, *θ_min_*, *θ_max_*, and *σ* are shared with Vm(θ). *C_Vf_* is a constant scaling factor to match the scale to the actual data. Then, the overall Vf was determined by weighting the magnitude of the angular deviation (W(Va)) to the contribution function of each leg (**Suppl. Fig. 4C**):

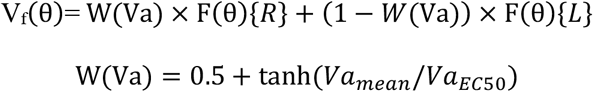

where *Va_mean_* is the mean value ofthe Va over past 100ms, *Va_EC50_* (200 °/s) determines the sensitivity of the weight to the Va value (Smaller *Va_EC50_* means that W(Va) reaches close to 1 at smaller Va).

We used the observed Va time series (actual data) for the input of the simulation. Therefore, the length of a simulation matched the length of a recording. We applied this simulation algorithm using each of 25 recording data to generate 50 leg-ipsilateral HS cell’s Vm and leg-contralateral HS cell’s Vm pairs, and Vf. Then, the simulated Vm tuning to the leg phase (**Suppl. Fig. 4E**), and simulated 2D behavioral maps (**Suppl. Fig. 4F**) were plotted using the simulated Vm and Vf time courses.

### Ball stop experiment

To compare the effect of the fly’s movement during walking vs. non-walking conditions (“Before”, “Ball Stopped”, “After” conditions), for each trial, we extracted a quarter of the total pixels whose intensity varied most over time from the side-view video recordings during BPNs activation. Then, the time course of the pixel intensity was high-pass filtered (> 5Hz) per pixel and averaged across extracted pixels. The autocorrelation coefficient of the filtered, averaged time course was calculated (**Suppl. Fig. 2A, bottom**). The periodicity strength of the fly’ movement (leg motion coupling) was estimated as the standard deviation of the autocorrelation coefficient trace over a window defined by 100–500ms lag. This was necessary since the coefficient of the autocorrelation at lag 0 was always 1 independent of the periodic nature of the pixels change in intensity, and therefore, the coefficient within a time window around lag 0 (0–100ms) were not used. Similarly, the periodicity strength of the corresponding neural activity (Vm coupling) was estimated as the standard deviation of the autocorrelation coefficient of Vm_|5Hz_over the 100–500ms lag time window. In **Suppl. Fig. 2B**, each data point represents the mean value of the Vm coupling over 6 trials per fly-cell pair, with bars indicating the grand mean. **Suppl. Fig. 2C**, shows for all the 6 trials from all fly-cell pairs (N=10) across the 3 conditions (6×10×3=180 data points) the Vm vs. leg motion couplings. Linear regression was performed for these 180 data points.

### External leg movement experiment

Using a side view image, the time points when the passive moving leg reached the maximal and minimal horizontal positions were extracted. Like the definition of leg phase during spontaneous walking and opto-runs, fictive stance and swing were defined as when their positions shifted from the local minima to the local maxima and vice versa, respectively. For **Fig. 4E**, the amplitude of the Vm oscillation within a stride window (from fictive stance onset to the next fictive stance onset) was calculated using raw Vm traces similar to **Fig. 3C**.

### Calcium imaging

XY motion was corrected using an algorithm described in ^101^. Relative fluorescence changes (ΔF/F) were calculated with respect to the baseline fluorescence during quiescence. Region of interests were defined as those pixels with mean intensities over a trial (120 s) 2 SD brighter than the average of all pixels. For calculating the cross-covariance and the walking velocity tuning (**Fig. 6F, G**), treadmill signals were down sampled to the imaging scanning rate (15 Hz). Based on the time lag in the cross-covariance (**Fig. 6F**, +666 ms), the walking velocity tuning was plotted with the lagged calcium response (Va bins: 40°/s vs. Vf bins: 1mm/s, **Fig. 6G**). The value at each pixel was calculated as the mean of all the collected data points. To extract walking and front leg grooming events (**Fig. 6H, I**), we annotated a fly’s side-view video using a deep learning-based algorithm, DeepEthogram ^102^. Considering the slow kinetics of calcium activity, walking or grooming events lasting over 500ms were selected, and the mean calcium response in each event was used for the analysis.

### Anatomy

A confocal image of the AN split-GAL4 driver was registered onto the JRC2018 unisex brain template ^103^ using the CMTK toolkit and then converted into a Color-Depth MIP image (**Suppl. Fig. 6C**) ^48^. A Color-Depth MIP image for the corresponding neuron in the hemibrain was generated using the Natverse toolkit ^104^. The shortest synaptic paths were analyzed on NeuPrint ^105^(**Suppl. Fig. 6E**).

### Statistics

We performed a two-sided Wilcoxon signed-rank test for paired groups, a two-sided Wilcoxon ranksum test for comparisons between two independent groups, and Kruskal-Wallis followed by a Tukey-Kramer test for multiple comparisons.

---

**Video S1, related to Fig. 3, Suppl. Fig. 1:** Example video showing the labels used to track two joints from the fly’s three legs of the left side. Color code indicates different labels: orange, hindleg femur-tibia joint; red, hindleg tibia-tarsus joint; light-green, middle leg femur-tibia joint; yellow middle leg tibia-tarsus joint; dark-blue, foreleg femur-tibia joint; light-blue, foreleg tibia-tarsus joint.

**Supplementary Fig. 1:**
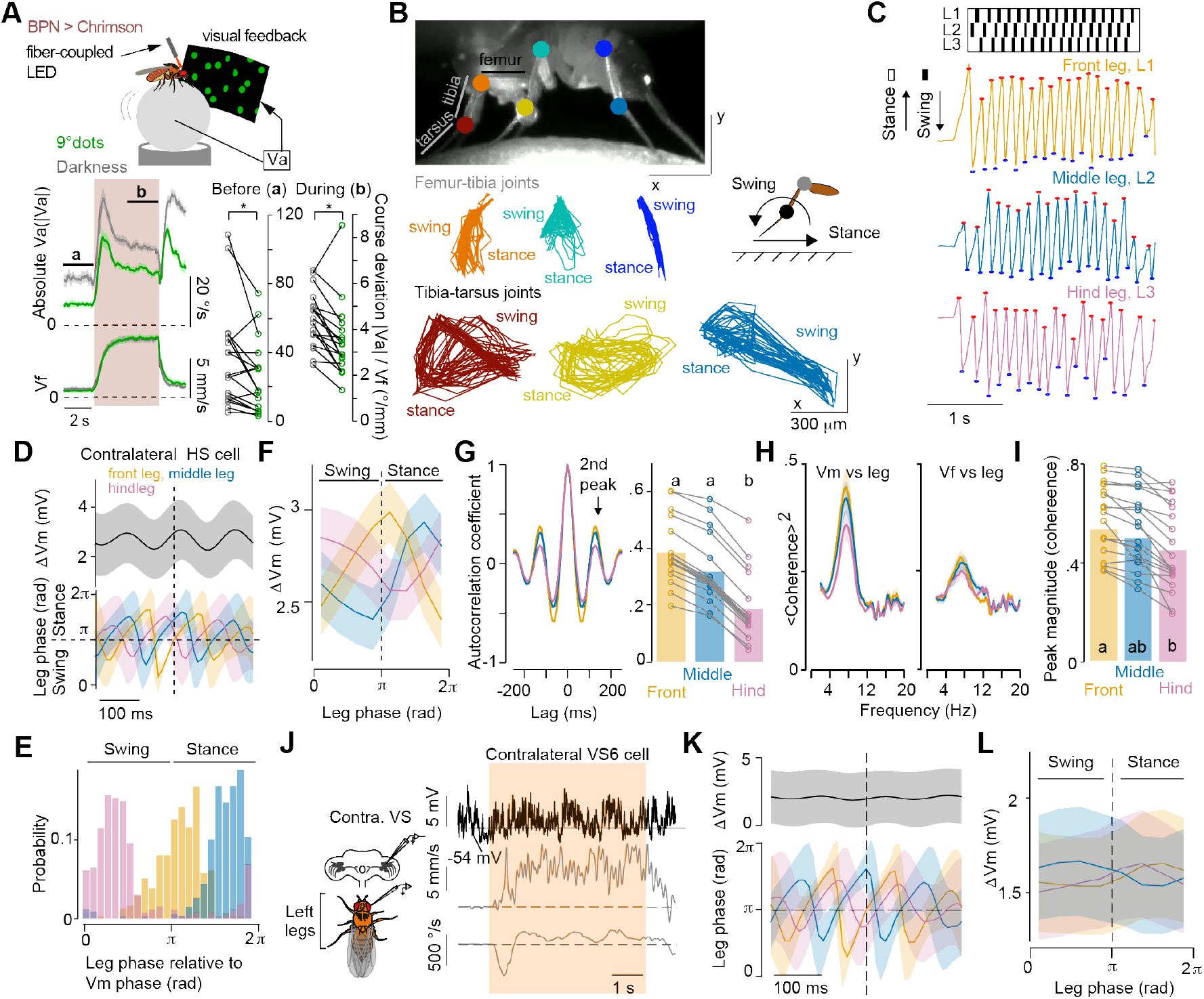
Leg movements and their correlation with neural activity and forward velocity. **(A)** Top, schematic of the closed-loop configuration between the fly’s angular velocity (Va) and the visual stimuli (“visual feedback”, green random dots of size 9°, see **Methods).** Bottom, the abosolute Va (top, |Va|) and the forward (middle, Vf) velocity of flies walking in darkness (gray) or in the presence of visual feedback (green, grand mean ± SEM, N=18 flies). The shaded area indicates the period of optogenetic activation of of BPNs, the black lines show the time used for analysis: before **(a)** and during **(b)** optogenetic stimulation. Right, mean course deviation, |Va|/Vf, before and during activation of BPNs in flies walking in darkness (black) or under the presence of visual feedback (green) (before: p=0.016, Z=2.42; during: p=0.022, Z=2.29, Wilcoxon signed-rank test). (B) Top, side view of an example fly with markers on the femur-tibia and tibia-tarsus joints labeled with DeepLabCut. Bottom, example trajectories of joints over the strides during a 5s activation of BPNs. Color code indicates the specific leg. (C) Top, extracted swing and stance phases of the three left legs. Bottom, corresponding time courses of the combined x-position of the femur-tibia and tibia-tarsus joints (see **Methods).** Red and blue markers indicate local maxima and minima of the position, corresponding to the onset of swing and stance, respectively. (D) Mean membrane potential change (ΔVm) and leg phases triggered at the stance onset of the left front leg (example fly, n=454 stance onsets). Shaded areas indicate SD. **(E)** Probability distribution of leg phases relative to the phase of Vm oscillations for the example in **(D).** Phase values from 0 to p and from π to 2π correspond to the swing and stance periods of the cycle, respectively. (F) HS cells tuning to the contralateral front (orange), middle (blue), and hind (pink) leg movements (grand mean ± SEM, N=19 fly-cell pairs). **(G)** Left, coefficient of autocorrelation of leg movements. Color code: same as in **(B).** Right, amplitude of the coefficient for the second peak in each leg. Letters indicate a significant difference (P<0.05, H=22.56, Kruskal-Wallis followed by Tukey Kramer test, N=19 flies). (H) Coherence between leg movements and Vm (left), and between leg movements and Vf (right) (N=19 fly-cell pairs). (I) Magnitude of the coherence peak between leg movement and Vm to each leg (P<0.05, H=6.32, Kruskal-Wallis followed by Tukey Kramer test, N=19 fly-cell pairs). (J) Left, schematic of the recorded neural activity and leg movements. Right, example time series of a right VS6 cell’s Vm (top), the fly’s Vf (middle) and Va (bottom) during activation in BPNs (orange shaded area). (K) Mean D Vm and leg phases triggered at the left front leg’s stance onset in the same example. Shaded areas indicate SD (n=775 stance onsets). (L) Tuning of contralateral VS cells’ Vm to the front (orange), middle (blue), and hind (magenta) leg phases (N=7 fly-cell pairs).

**Supplementary Fig.2:**
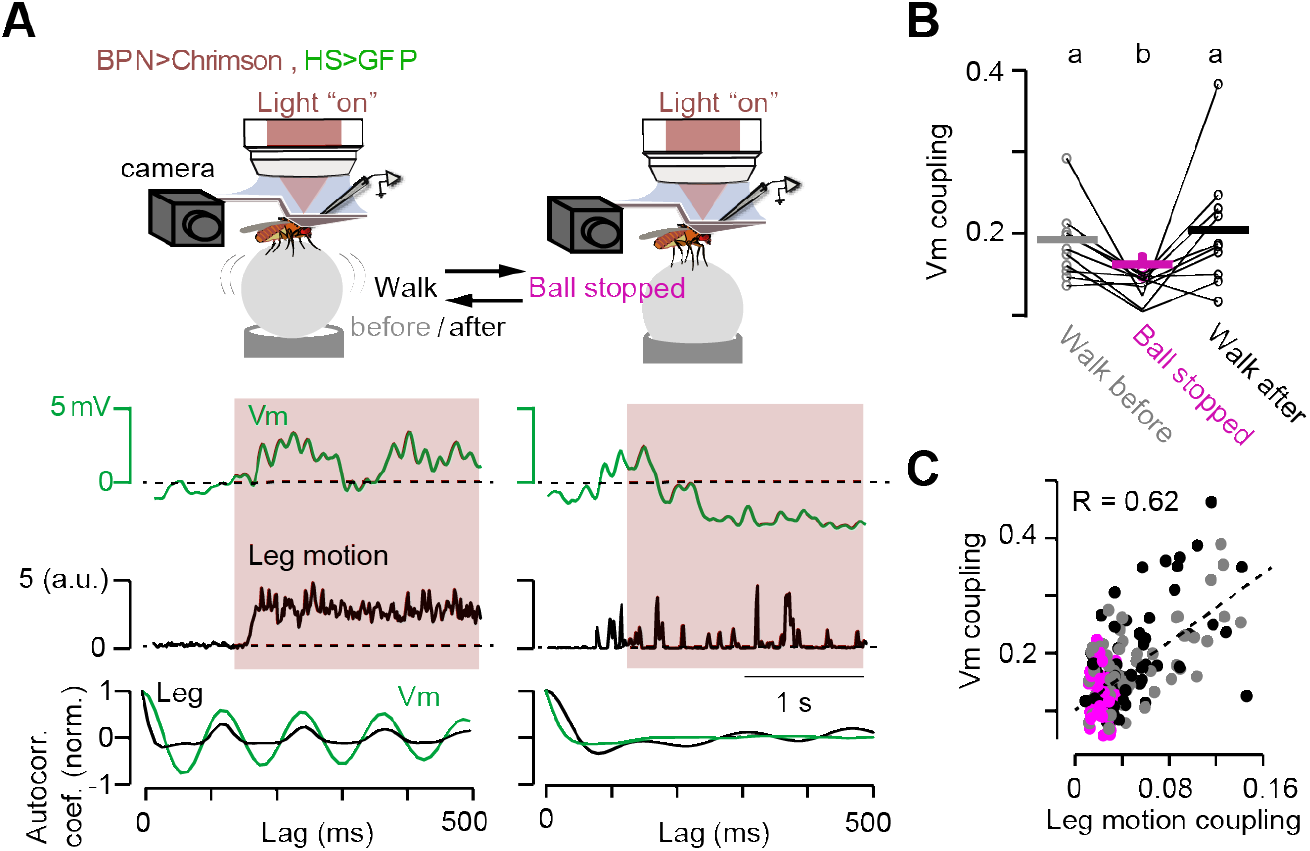
Activation of BPNs *per se* does not induce oscillations in the activity dynamics of HS cells. **(A)** Top, schematic of the experimental design. Whole cell-patch recordings from HS cells in simultaneous with optogenetic activation of BPNs (red shadow, “lights on”) in walking (“walk”, left) or not-walking (“ball stopped”, right) flies. The latter condition was induced by stopping the air flow of the ball; note that this manipulation was reversible. To monitor behavior under these conditions, overall leg motion was extracted from the camera tracking legs (see **Methods).** Middle, time series of the membrane potential (Vm) and the combined leg motion signal. Bottom, normalized autocorrelation coefficient as a function of lag for the neural activity dynamics (Vm, green) and the leg motion signal (leg, black). The oscillatory profile of the autocorrelation for the leg motion signal is characteristic of the periodic nature of leg movement during walking. (B) Quantification of the neural activity autocorrelation strength (Vm coupling) before, during, and after stopping the ball. Letters above the plot indicate significant differences (p<0.05, Kruskal-Wallis followed by Tukey Kramer test, Z=11.34, N=10 fly-cell pairs). **(C)** Correlation between the autocorrelation strength of the fly’s leg movements (leg motion coupling) and Vm coupling in each trial. The dashed line shows the linear regression of all the points (n=180 trials).

**Supplementary Fig.3:**
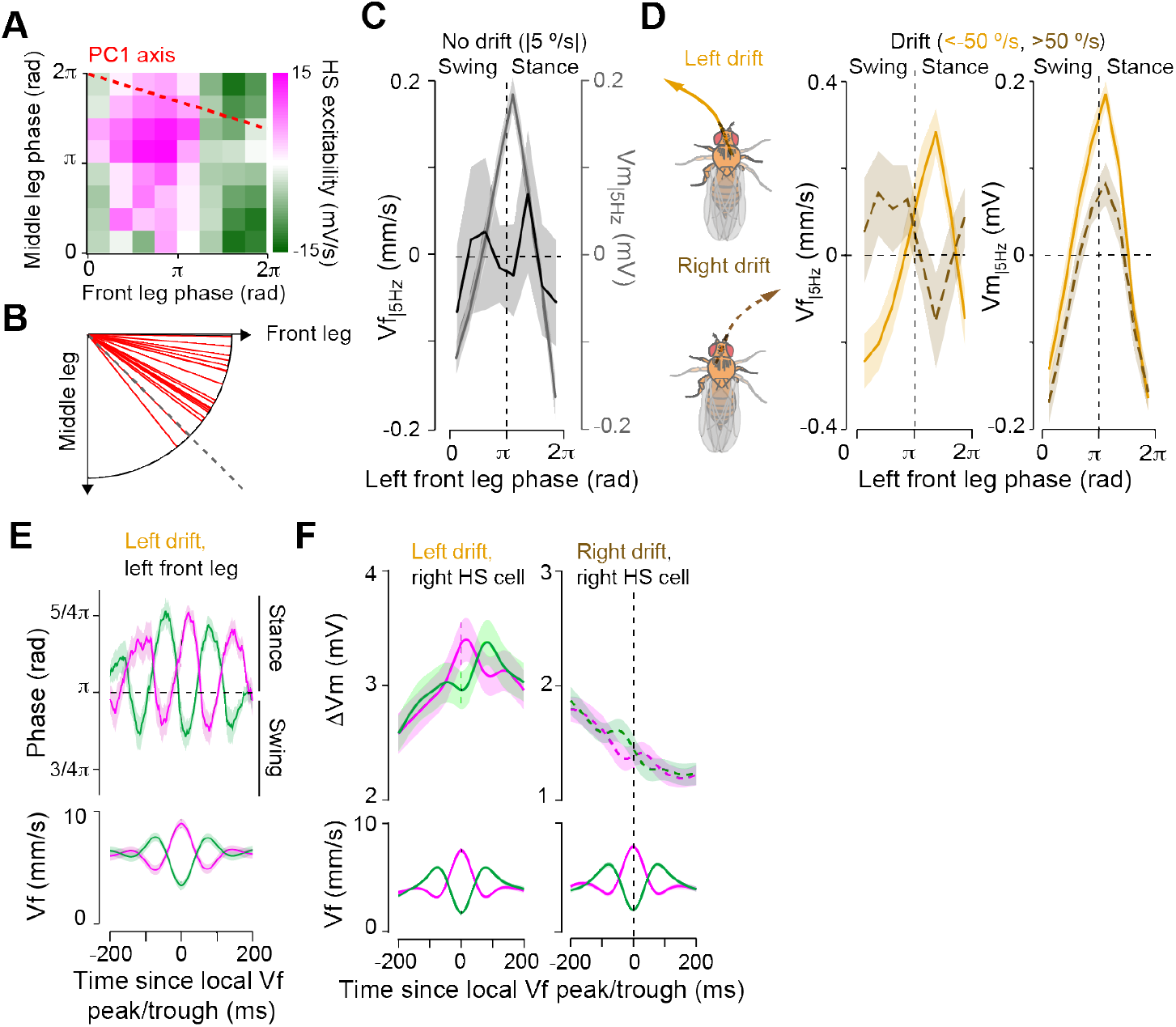
The relation between the stride-cpupled fluctuations in Vf and oscillations in HS cells depend on the direction of drift from straight course. **(A)** HS cells excitability (see **Methods)** during opto-runs projected onto a 2D leg phase space; x-axis: left front leg, y-axis: left middle leg. The PC 1 axis of the 2D map, the orientation of the largest gradient, is indicated by the red, dashed line. Data were collected from 19 fly-cell pairs. (B) PCI phase axis calculated per fly-cell pair (N=19 fly-cell pairs). The dashed line represents the alignment of the PC 1 axis to both front and middle legs. (C) Tuning of Vf (high-pass filtered, Vf_|5Hz_) and right HS cells (high-pass filtered, Vm_|5Hz_) to the stride cycle of the left front leg during walking with marginal angular drift (−5<V_a_<5°/s; N=19 fly-cell pairs, grand mean ± SEM). The double peak of Vf over the stride cycle indicates a symmetric contribution to the fly’s speed by the right-left pairs of legs. **(D)** Same as in **(C)** but for opto-runs with left (Va<−50°/s, solid orange curve) or right (Va>50°/s, dashed maroon curve) angular drift (N=18 fly-cell pairs, grand mean ± SEM). The stronger contribution to Vf by the leg ipsilateral to the drift direction dominates the peak in Vf over the stride cycle. In contrast, the peak of Vm over the stride cycle does not change with angular drift. (E) Left front leg phase (top) and Vf (bottom) triggered at the local peak (magenta) or trough (green) of the Vf in walking segments with left angular drift (N=18 fly-cell pairs, grand mean ± SEM). (F) Change in the activity of HS cells (ΔVm, top) and Vf (bottom) triggered at the local peak (magenta) or trough (green) of the Vf_|5Hz_ spontaneous walking segments with left (N=23 fly-cell pairs, left) or right (N=24 fly-cell pairs, right) angular drift. Traces indicate grand mean ± SEM.

**Supplementary Fig.4:**
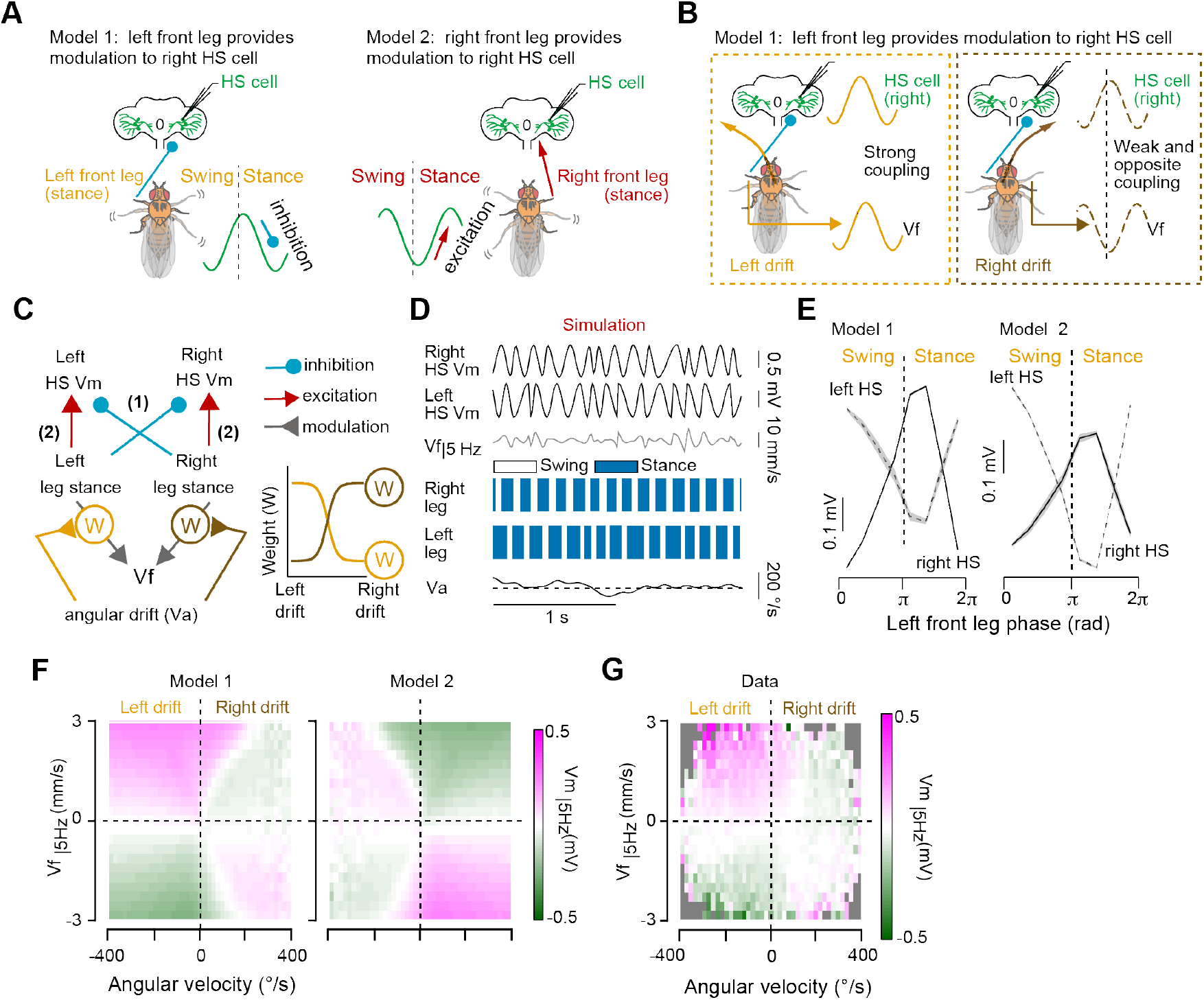
A simple model supports the contribution of the contralateral front leg to the stride-coupled modulation of an HS cell. **(A)** Two models with different origin for the stride-coupled modulation in HS cells. In model 1, the contralateral front leg drives an inhibitory signal during stance, whereas model 2 proposes an excitatory drive from the ipsilateral front leg during the stance phase. (B) Schematics showing the relationship between the right HS cell’s Vm and Vf under model 1. (C) Schematic of the circuit diagram for the simulation. Both right and left legs contributed to Vf with a weight (W) that was proportional to the ongoing Va, i.e., the direction of the fly’s angular drift. (D) Example of a simulated time course for the right (1st row) and left (2nd row) HS cell’s Vm, Vf_|5Hz_ (3rd row), right (4th row) and left (5th row) leg phases, and angular velocity (Va) (6th row). (E) Simulated tuning of right (solid line) and left (dashed line) HS cells under models 1 (left) and 2 (right). N=50 simulated cells. (*F*) Simulated activity of right HS cells (high-pass filtered, Vm_|5Hz_) as a function of the angular (Va) and forward (high-pass filtered, Vf_|5Hz_) velocities of the fly based on model 1 (left) and model 2 (right). Simulations were performed with 50 cells. (G) Activity in right HS cells (high-pass filtered, Vm_|5Hz_, N=25 fly-cell pairs) as a function of the angular (Va) and forward (high-pass filtered, Vf_|5Hz_) velocities of the fly.

**Supplementary Fig.5:**
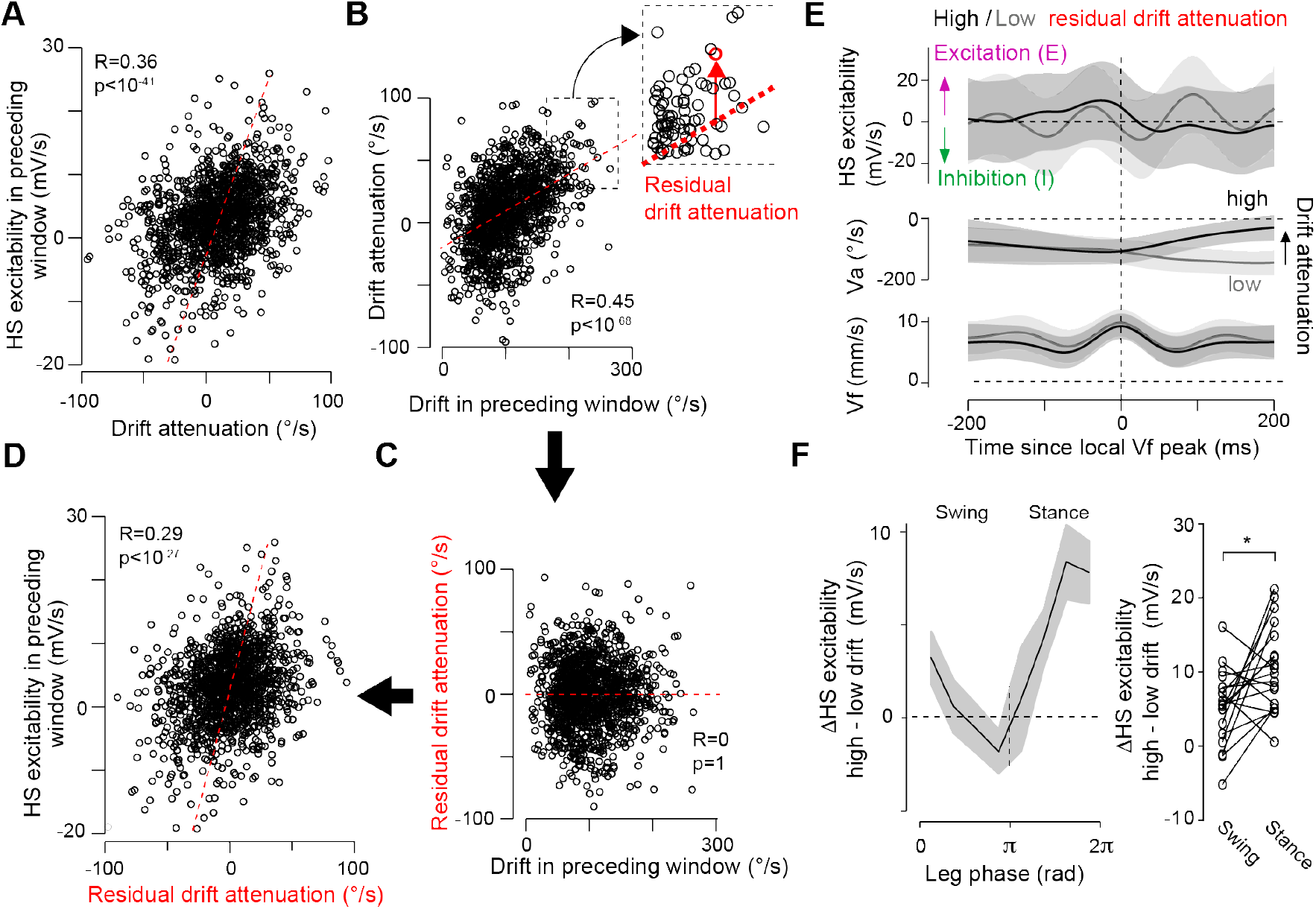
The excitability of HS cells over a stride strongly correlates with rapid steering independent of the state of Va. **(A).** Mean HS cells’ excitability per opto-run segment before the peak of Vf (time window: 200ms) vs. mean drift attenuation in the following 200ms. The dotted lines indicate the linear regression (R=0.36, p<10^-41^, n=1378 segments from 19 fly-cell pairs). (B) Mean drift attenuation per opto-run segment over 200ms after the local peak of Vf vs. mean angular drift per segment in the preceding 200ms (n=1378 segments from 19 fly-cell pairs). The dotted lines indicate the linear regression (R=0.45, p<10^-68^) Inlet: definition of “residual drift attenuation” for an example data point (see also **Methods). (C)** Mean residual drift attenuation per opto-run segment vs. the mean preceding angular drift over 200ms before the local peak of Vf. The dotted lines indicate the linear regression (by definition, R=0, p=1). (D) Mean HS cells’ excitability per opto-run segment over 200ms before the local peak of Vf vs. mean residual drift attenuation in the following 200ms. The dotted lines indicate the linear regression (R=0.29, p<10^-27^). (E) HS cells’ excitability (top), Va (middle) and Vf (bottom) triggered at the local peak of Vf, in opto-run segments with similar magnitude of angular drift before the local peak of Vf (leftward direction, Va<−50°/s), but with low (gray) or high (black) residual drift attenuation 200ms after the local peak of Vf (n=360 segments; mean ± SD, segments were collected from 19 fly-cell pairs). (F) The difference in HS cell’s excitability over a stride cycle of opto-run segments with low (50°/s<Va<~0°/s) vs. high (200°/s<Va<−150°/s) angular drifts (see **Fig. 5E).** The trace and shaded area represent the grand mean ± SEM, respectively (p=0.020, Z=−2.33, Wilcoxon signed-rank test, N=19 fly-cell pairs).

**Supplementary Fig.6:**
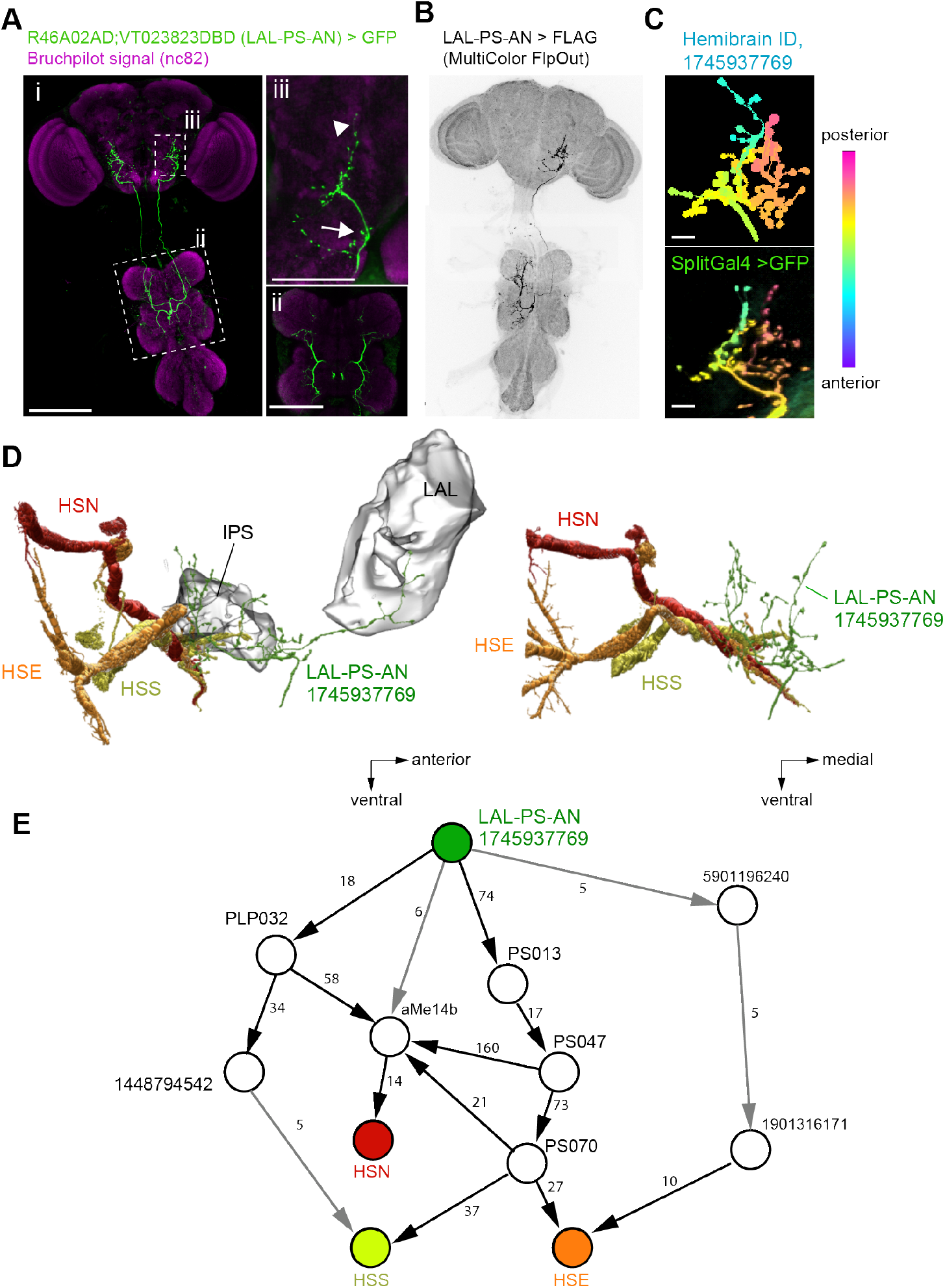
Anatomy of LAL-PS-AN_contra_, a class of ascending neurons projecting to IPS and LAL. **(A)** Projected z-stacked confocal image of the split-GAL4 line R46A02AD-VT023823DBD driving expression of EGFP (i). This line labels a class of ascending neurons that innervates leg neuropil within the proto- and mesomeres of the VNC (ii), and with brain projection fields within the GNG, IPS (arrow), and LAL (arrowhead) regions (iii). (B) A multi-color flip-out image revealing the contralateral projections of an individual LAL-PS-AN_contra_ (C) Maximum intensity projection images (MIPs) of an aligned confocal image of our split-GAL4 (bottom) and a putative corresponding neuron identified in the hemibrain EM dataset. Color represents different frames of the image stack in the antero-posterior axis. (D) EM-based reconstructed HS cells and the putative LAL-PS-AN_contra_ neuron (green) from the hemibrain dataset (see **Methods). (E)** Shortest possible path between the putative LAL-PS-AN_contra_ neuron and HS cells. Synaptic weights: ≥5 synaptic contacts, gray; ≥10 synaptic contacts, black. Scale bars: 100 μm **(A, B);** 10μm **(C).**

**Supplementary Fig.7:**
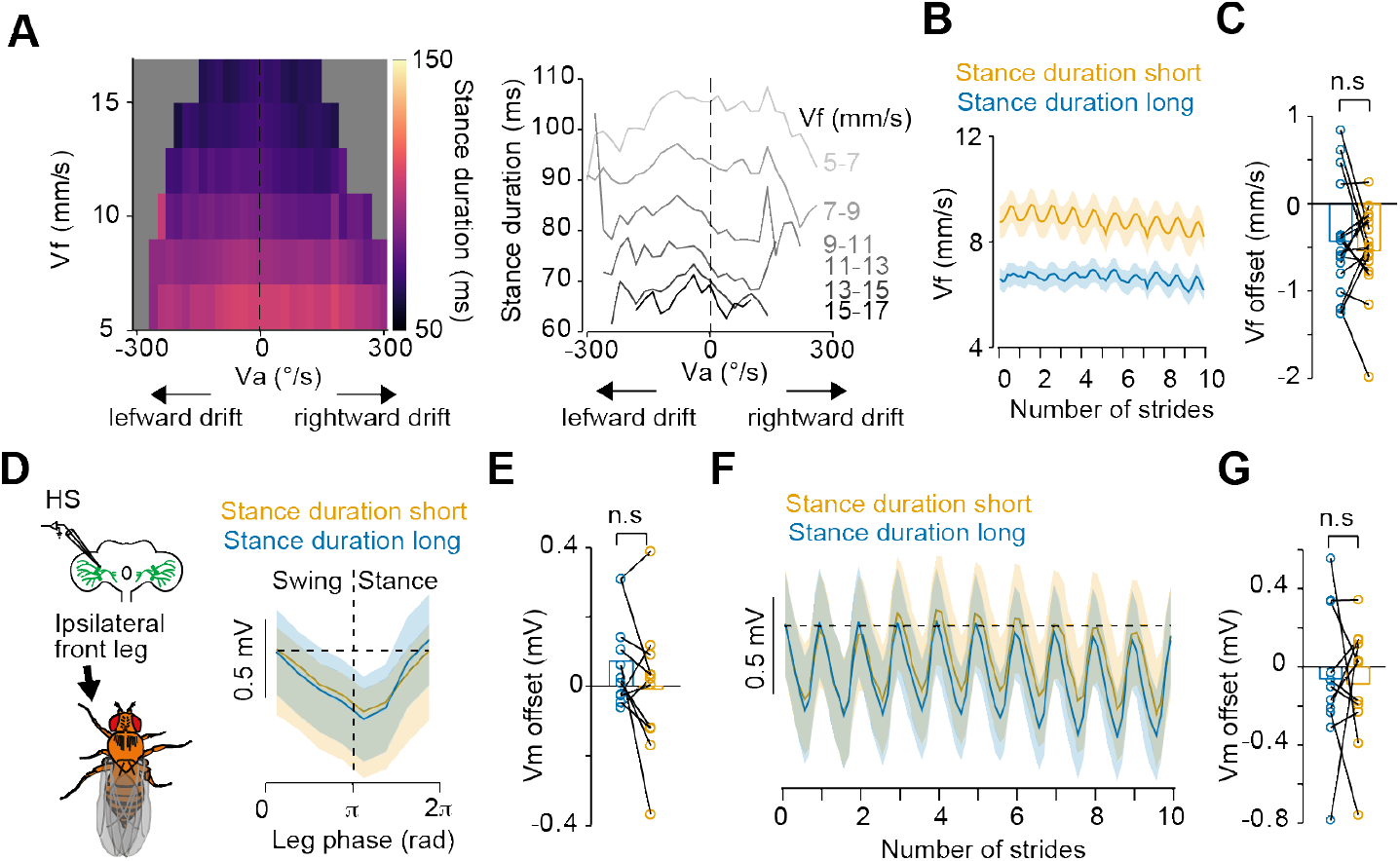
Relationship among the stance duration, forward velocity and left HS cell activity. **(A)** Left, front leg stance duration as a function of the angular (x-axis, Va, 20 °/s bins) and forward (y-axis, Vf, 2 mm/s bins) velocities of the fly. Right, the mean stance duration as a function ofVa at different Vf. Data were collected from 19 flies. (B) Vf tuning to front leg stride cycle over ten strides for shorter (orange) or longer (blue) stance duration (grand mean ± SEM, N=17 flies). **(C)** The Vf offset at the end of ten strides relative to the beginning for shorter and longer stance durations (p=0.62, Z=0.50, N=17 flies, Wilcoxon signed-rank test). (D) Left, schematic of the experimental configuration. Right, left HS cells tuning to the left front leg phase for shorter (orange) or longer (blue) stance durations (grand mean ± SEM, N=11 fly-cell pairs). **(E)** The Vm offset at the end relative to the beginning of the stride for shorter and longer stance durations (N=11 fly-cell pairs, p=0.28, Wilcoxon signed-rank test). **(F)** Same as **(E),** but traces over ten strides. **(G)** Same as in **(E)** but for ten consecutive strides (p=0.97, N=11 fly-cell pairs, Wilcoxon signed-rank test).

## Notes

### Competing Interest Statement

The authors have declared no competing interest.

